# Adaptor Protein-3 Produces Synaptic Vesicles that Release Phasic Dopamine

**DOI:** 10.1101/2023.08.07.552338

**Authors:** Shweta Jain, Andrew G. Yee, James Maas, Sarah Gierok, Hongfei Xu, Jasmine Stansil, Jacob Eriksen, Alexandra Nelson, Katlin Silm, Christopher P. Ford, Robert H. Edwards

## Abstract

The burst firing of midbrain dopamine neurons releases a phasic dopamine signal that mediates reinforcement learning. At many synapses, however, high firing rates deplete synaptic vesicles (SVs), resulting in synaptic depression that limits release. What accounts for the increased release of dopamine by stimulation at high frequency? We find that adaptor protein-3 (AP-3) and its coat protein VPS41 promote axonal dopamine release by targeting vesicular monoamine transporter VMAT2 to the axon rather than dendrites. AP-3 and VPS41 also produce SVs that respond preferentially to high frequency stimulation, independent of their role in axonal polarity. In addition, conditional inactivation of VPS41 in dopamine neurons impairs reinforcement learning, and this involves a defect in the frequency dependence of release rather than the amount of dopamine released. Thus, AP-3 and VPS41 promote the axonal polarity of dopamine release but enable learning by producing a novel population of SVs tuned specifically to high firing frequency that confers the phasic release of dopamine.

**Significance statement:** Reinforcement learning requires the phasic dopamine produced by burst firing but synaptic vesicle depletion limits the ability to convey information at high firing rates. We now find that AP-3 has two independent roles in dopamine release. First, AP-3 confers the axonal polarity of dopamine release by targeting vesicular monoamine transporter 2 (VMAT2) to the axon. Second, AP-3 acting locally at the nerve terminal produces synaptic vesicles that respond specifically to high frequency stimulation. Consistent with this, loss of AP-3 impairs reinforcement learning and this reflects the defect in release at high frequency, not the reduction in axonal dopamine.

## Introduction

The nervous system encodes information through the timing and frequency of action potentials. The speed of synaptic transmission conveys information about timing, but at many synapses, repeated stimulation depletes synaptic vesicles, limiting release at high frequency. In contrast, the release of monoamines and other neuromodulators shows a steep dependence on firing rate. Tonic release of dopamine reflects baseline firing at 3-8 Hz (1) whereas phasic release responds specifically to high frequency firing (25-40 Hz or more), and phasic dopamine signals the positive reward prediction error required for reinforcement learning (2, 3). Independent of burst firing in the cell body, dopamine release ramps up in anticipation of reward due to acetylcholine acting locally in the striatum (4–6). However, the mechanism responsible for increased dopamine release with strong stimulation remains unknown. Dopamine also differs from most other classical transmitters in the sites of release, with somatodendritic release in the midbrain as well as axonal release in the striatum (7–11). The mechanisms responsible for the polarity of dopamine release and the physiological role of somatodendritic release are poorly understood. The synaptic vesicles (SVs) that store monoamines appear to differ from those that contain other classical transmitters (12, 13), suggesting that these differences may underlie the distinct frequency dependence and polarity of dopamine release.

SVs form by local, endocytic recycling at the nerve terminal. Clathrin and its heterotetrameric adaptor protein AP-2 regenerate SVs either directly from the plasma membrane (14) or immediately after an ultrafast form of endocytosis (15–17). A defect in this recycling mechanism impairs the response to stimulus trains due to SV depletion (18–20). Originally identified through work on endolysosomal trafficking in yeast (21), the related adaptor AP-3 also produces SVs, but AP-3 produces SVs from endosomes rather than the plasma membrane (22–24). In *C. elegans*, AP-3 promotes the axonal trafficking required for assembly of presynaptic release machinery (25, 26). In mammals, loss of AP-3 causes seizures, reduces the number of SVs (27) and influences SV composition (28–31). However, mammalian nerve terminals form without AP-3 (32) and the analysis of glutamatergic transmission has revealed only a modest change in asynchronous release and synaptic depression (30, 33-36). The physiological consequences of a role for AP-3 in SV biogenesis have thus remained uncertain.

The analysis of neurotransmitter corelease now suggests a physiological role for AP-3. A subset of midbrain dopamine neurons release glutamate as well as dopamine (37–40). These populations express both vesicular monoamine transporter VMAT2 and vesicular glutamate transporter VGLUT2 (41). Despite expression in the same neurons, VMAT2 and VGLUT2 segregate at least in part to different microdomains (12) and differ in response to stimulation (13). In previous work, we found that loss of AP-3 preferentially impairs the regulated exocytosis of vesicular monoamine transporter VMAT2^+^ rather than VGLUT2^+^ vesicles (13), suggesting a specific role in monoamine storage and release. We now find that AP-3 is required for the axonal polarity of dopamine release. We also find that, independent of its role in polarity, AP-3 produces a subpopulation of SVs that respond preferentially to stimulation at high frequency, and dopamine release from AP-3-derived SVs contributes to reinforcement learning. Rather than a role defined by transmitter, AP-3 thus contributes specifically to release at high frequency.

## Results

### AP-3 Confers the Polarity of Dopamine Storage

AP-3 is a heterotetrameric complex that occurs in two forms, a ubiquitous form that directs trafficking of membrane proteins to lysosomes and lysosome-related organelles (42) and a form specific to neuroendocrine cells which is less well understood. The large δ subunit is common to both forms but two other subunits (β and µ) occur in two isoforms, one ubiquitous and the other neural, that define the ubiquitous and neural isoforms of AP-3. To determine whether the loss of striatal dopamine reflects the well-established role for AP-3 in trafficking to lysosomes, or a more specific defect in trafficking to neurosecretory vesicles, we thus analyzed tissue dopamine content in mice lacking either the ubiquitous β subunit β3A (43) or the neural subunit β3B (23). Very similar to the complete loss of AP-3 (13), loss of the neural subunit β3B reduces striatal dopamine by ∼50% whereas loss of the ubiquitous β3A has no effect (Fig. 1A). Remarkably, we now find that loss of β3B increases midbrain dopamine content (Fig. 1B), an effect also observed in *mocha* mice lacking the δ subunit (*SI Appendix*, Fig. S1A), and β3A again has no effect. AP-3 thus regulates the balance between axonal and somatodendritic dopamine stores. Surprisingly, the β3A knockout (KO) reduces striatal as well as midbrain levels of serotonin (*SI Appendix*, Fig. S1B), indicating that the ubiquitous form of AP-3 has a larger role in serotonergic than dopaminergic neurons.

**Fig. 1.**
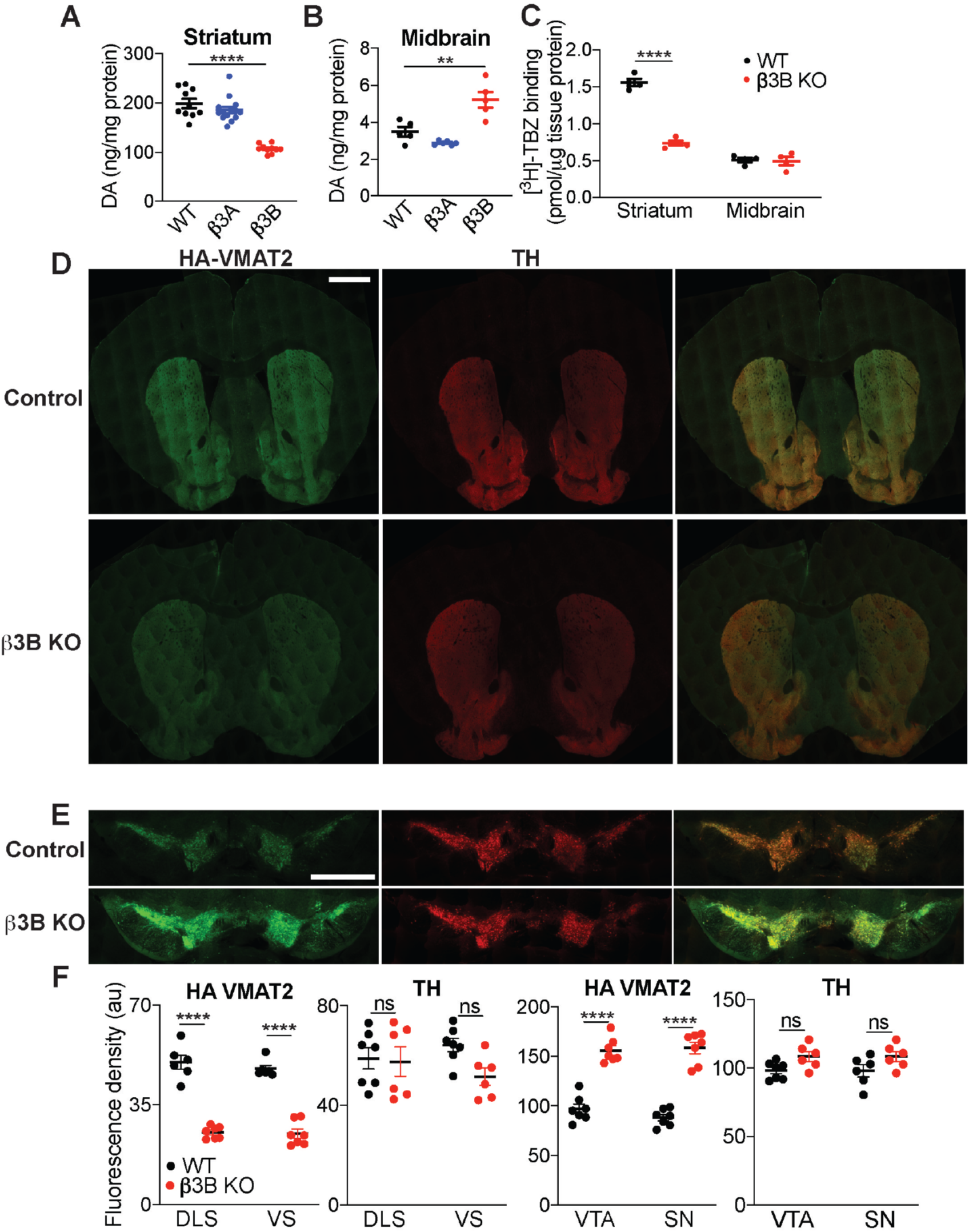
Loss of neural AP-3 disrupts the polarity of monoamine storage *in vivo*. (A) Scatterplot of tissue dopamine (DA) content in the striatum of WT, β3A and β3B KO mice. ****, p<0.0001 by one-way ANOVA with *post hoc* Bonferroni’s test (n=10 for WT, β3B KO, n=14 for β3A KO; WT mice were littermates derived from heterozygous β3B parents and all animals are on a C57BL/6 background. (B) Tissue content of DA in the midbrain of WT, β3A, and β3B KO mice. **, p=0.0021 by one-way ANOVA with Bonferroni’s multiple comparisons test (n=5 for WT and β3B KO, n=6 for β3A KO). (C) Specific ^3^H-tetrabenazine binding in the striatum and midbrain of WT and β3B KO mice. ****, p<0.0001 by two-way ANOVA with Sidak’s multiple comparison test (n=4 for WT and β3B KO). (D,E) Striatal (D) and midbrain (E) slices from WT and β3B KO/HA-VMAT2 BAC transgenic mice were immunostained for HA (green) and TH (red). (F) Mean fluorescence per μm^2^ area from individual fields such as (D) and (E). DLS, dorsolateral striatum, VS, ventral striatum, VTA, ventral tegmental area, SN, substantia nigra. ****, p<0.0001; ***, p<0.001; ns, not significant by two-way ANOVA with Sidak’s multiple comparisons test (n=6 fields from three mice for each condition). Scale bars, 1 mm. Error bars indicate SEM.

The subcellular localization of vesicular monoamine transporter VMAT2 defines the vesicles capable of monoamine storage and release. To determine whether AP-3 influences the distribution of this transporter *in vivo*, we first used binding to the VMAT2 inhibitor tetrabenazine (TBZ). Figure 1C shows that loss of β3B reduces striatal TBZ binding by >50%. The KO has no effect on TBZ binding in the midbrain, but axonal projections from serotonergic neurons to the midbrain complicate the interpretation of midbrain VMAT2 levels. We thus immunostained sections from HA-VMAT2 BAC transgenic mice (13) with and without β3B, confirming the reduction in striatal VMAT2 within the axonal projection of dopamine neurons labeled for the catecholamine biosynthetic enzyme tyrosine hydroxylase (TH) (Figures 1D,F and *SI Appendix*, Fig. S1D). HA immunoreactivity increases within the TH^+^ cell bodies and dendrites of midbrain dopamine neurons (Fig. 1E,F and *SI Appendix*, Fig. S1E). However, the striatal staining for TH and the dopamine transporter (DAT) does not change significantly in overall density (*SI Appendix*, Fig. S1C) or fluorescence per process length (*SI Appendix*, Fig. S1D and E). The shift in distribution of dopamine stores observed in the absence of neuronal AP-3 thus reflects a normal role for the adaptor in membrane trafficking of VMAT2 to the axon rather than dendrite. Consistent with previous work in *C. elegans* (25, 26), mammalian AP-3 thus plays a role in axonal polarity, specifically in the storage of monoamine.

To understand how AP-3 contributes to polarity, we determined the location of the two adaptor isoforms. Since loss of any subunit destabilizes the AP-3 complex, we took advantage of isoform-specific knockouts and a monoclonal antibody to the common δ subunit. We used hippocampal neurons because midbrain cultures require a glial monolayer expressing substantial amounts of AP-3 that complicate analysis of the cocultured neurons, and VMAT2 trafficks similarly in hippocampal and midbrain dopamine neurons (13, 44). In wild-type mouse neurons, the δ antibody labels both dendrites (identified by double staining for MAP2) and axons (identified by double staining for the initial segment protein ankyrin G) (*SI Appendix*, Fig. S2). Loss of β3A reduces but does not eliminate the δ immunoreactivity in both MAP2^+^ and MAP2^-^ processes (*SI Appendix*, Fig. S2 A,B). Loss of β3B also leaves residual staining (i.e., β3A) in the dendrites but essentially eliminates δ immunoreactivity in the axon (*SI Appendix*, Fig. S2). Thus, both isoforms of AP-3 localize to dendrites but the neural isoform predominates in the axon. Somatodendritic localization of neural AP-3 presumably contributes to the polarity of dopamine storage, targeting VMAT2 and other membrane proteins to the axon. Axonal localization of the neural isoform supports a role in SV recycling and biogenesis (22, 23).

### AP-3 and VPS41 Confer the Frequency-Dependent Exocytosis of Monoamine SVs

AP-3 targets VMAT2 to axonal SVs but the distinct properties of dopamine release (13) suggested that the adaptor might have a larger role in the acquisition of specific release properties. To test this possibility, we imaged the externalization and recycling of VMAT2 using a lumenal fusion to the pH-sensitive pHluorin variant of GFP (VMAT2-pH), which shows quenching at the low pH of SVs and an increase in fluorescence with exposure to higher pH at exocytosis; the reacidification that follows endocytosis results in requenching (44). Introduced into midbrain neurons identified as dopaminergic using a fluorescent ligand for the dopamine transporter (45) (Figure 2A), VMAT2-pH shows ∼70% reduction in the peak response to 10 Hz stimulation in cells from the β3B KO relative to wild-type and the β3A KO (Fig. 2B), very similar to the complete loss of AP-3 in *mocha* neurons (13). A similar fusion to the vesicular glutamate transporter VGLUT2 (VGLUT2-pH) shows a much smaller reduction in release in hippocampal neurons from the β3B KO (Fig. 2C). However, the role of neural AP-3 in trafficking of VMAT2 is not specific to midbrain dopamine neurons. Expression of VMAT2-pH in hippocampal neurons from both β3B knockout and *mocha* mice shows a similarly reduced response relative to wild-type (*SI Appendix*, Fig. S3). Thus, the neural isoform of AP-3 is required for the presynaptic trafficking of VMAT2 in hippocampal as well as midbrain neurons, and the differences in trafficking from VGLUT2 do not reflect a specific role for AP-3 in only monoamine neurons. The preferential effect on VMAT2 suggests that AP-3 produces a distinct subpopulation of SVs enriched in VMAT2 but not VGLUT2. AP-3 associates with a number of other trafficking proteins including a subunit of the homotypic fusion and protein sorting (HOPS) complex, VPS41 (46, 47). Among these, VPS41 is particularly important for the formation of neurosecretory vesicles (48). VPS41 contains a clathrin heavy chain repeat and reversibly assembles into a lattice, suggesting it may function as the coat protein for AP-3. We now find that midbrain dopamine neurons from VPS41 conditional KO (cKO) mice also show an impaired response to stimulation (Fig. 2C). Thus, VPS41 may cooperate with AP-3 in neurons as well as endocrine cells. It is important to note that dense core vesicles invariably appear as isolated, discrete exocytic events when imaging pHluorin fusions to neural peptides (48–50). The graded response of VMAT2-pH to stimulation thus indicates localization to SVs rather than dense core vesicles.

**Fig. 2.**
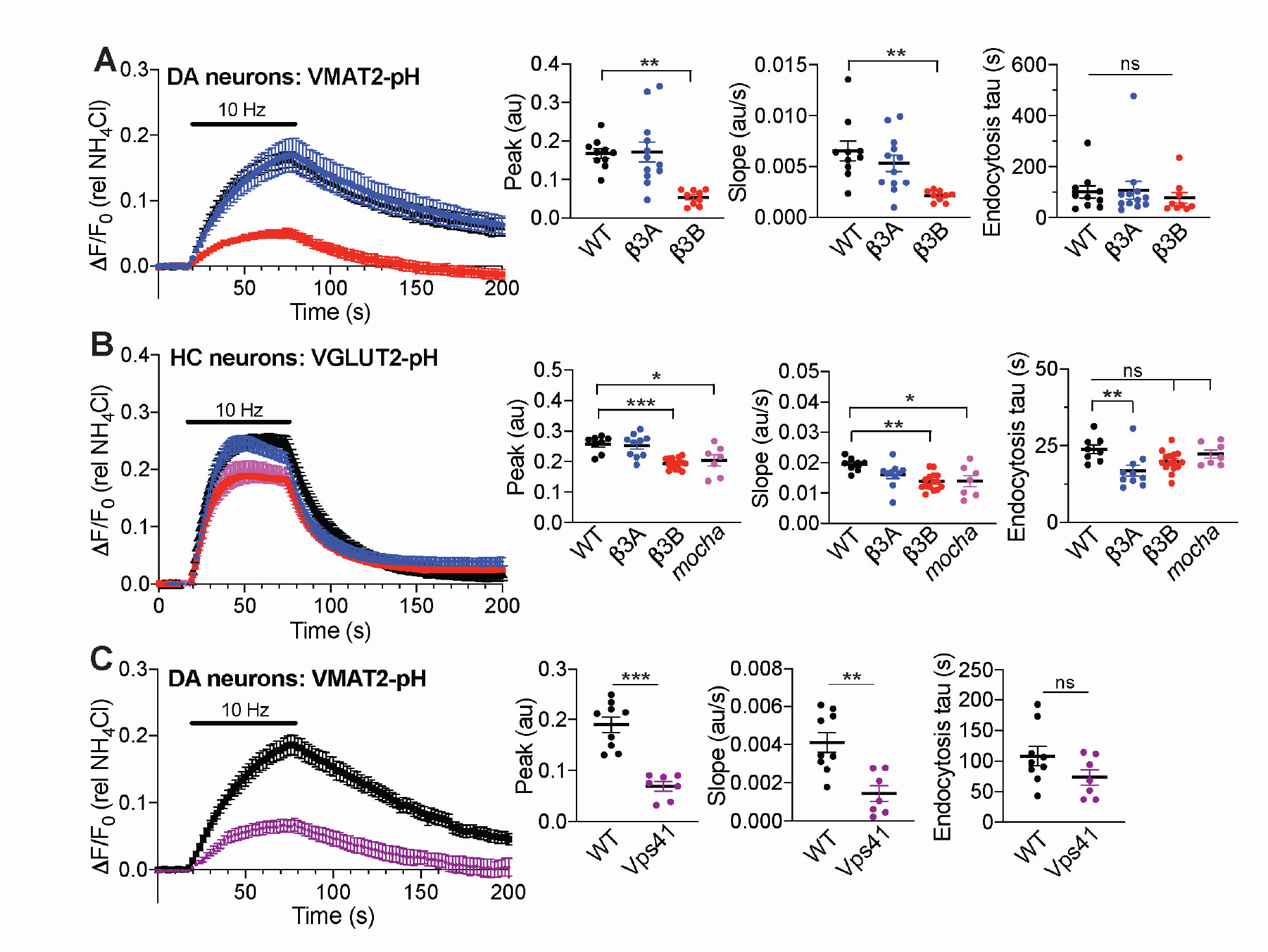
Loss of AP-3 impairs the regulated exocytosis of VMAT2^+^ vesicles to a greater extent than that of VGLUT2^+^ vesicles. (A) Response of VMAT2-pHluorin (VMAT2-pH) to stimulation at 10 Hz for 60 s in the midbrain dopamine neurons (identified with DAT ligand JHC-164) of WT, β3A and β3B KO mice. Scatterplots (right) indicate the mean peak response, mean initial slope at stimulation, and mean endocytic time constant. **, p<0.01; ns, not significant by one-way ANOVA with *post hoc* Bonferroni’s test (n=10 coverslips for WT, 12 for β3A KO and 9 for β3B KO from 3-4 different cultures). (B) Response of VGLUT2-pH to stimulation at 10 Hz for 60 seconds in the hippocampal neurons of WT, β3A KO, β3B KO and *mocha* mice, with the data analyzed as in (A). *, p<0.05; **, p<0.01; ***, p<0.001; ns, non-significant by one-way ANOVA with *post hoc* Bonferroni’s test (n=8 coverslips for WT, 10 for β3A KO, 15 for β3B KO and 7 for *mocha* from 3-4 different cultures). (C) Response of VMAT2-pH upon stimulation at 10 Hz for 60 seconds in midbrain dopamine neurons of WT and Vps41 KO mice, analyzed as in (A). **, p<0.01; ***, p<0.001; ns, not significant by Mann-Whitney test. n=9 coverslips for WT, 7 for Vps41 KO from 3 different cultures. The fluorescence was normalized to alkalinization in 50 mM NH_4_Cl.

Loss of VPS41 also reduces dopamine levels in the striatum, very similar to loss of AP-3 although midbrain dopamine does not increase significantly (*SI Appendix*, Fig. S4 A and B). Consistent with inactivation of VPS41 specifically in dopamine neurons, the VPS41 cKO does not affect serotonin levels in either striatum or midbrain. Loss of VPS41 also does not reduce axonal TH, arguing against a role for degeneration in the phenotype (*SI Appendix*, Fig. S4C) which has been suggested by previous work (51). The phenotype of VPS41 inactivation thus resembles that of AP-3.

In previous work, we found that VMAT2-pH shows a more linear response to the frequency of stimulation than VGLUT2-pH, which shows depression at high frequency (13). Although the role of AP-3 in axonal polarity of VMAT2 trafficking accounts for the reduced striatal dopamine stores, it cannot account for the reduced exocytosis of VMAT2^+^ vesicles because the defect in exocytic response occurs despite normalization to the reduced axonal VMAT2 revealed by alkalinization in NH_4_Cl (Fig. 2A,C, *SI Appendix*, Fig. S3). An independent role for AP-3 in SV regeneration might therefore account for the response of VMAT2^+^ SVs to stimulation at high frequency. Thus, we stimulated midbrain dopamine neurons expressing VMAT2-pH at either 5, 10, 25 or 50 Hz; we used the H^+^ pump inhibitor bafilomycin to block reacidification and hence focus on the exocytic response. Under these conditions, the response of VMAT2-pH to stimulation at 5 or 10 Hz does not differ significantly from wild-type in the proportion of SVs available for release (pool size), but the rate of initial response is reduced (Fig. 3A). At 25 or 50 Hz, however, the loss of neural AP-3 substantially reduces both pool size and the rate of response (Fig. 3A). The change in initial rate indicates that although the stimulation is prolonged, the role of neural AP-3 emerges within one second, suggesting relevance for the dopamine released *in vivo* by a short burst and arguing against a role for SV regeneration in the observation; even ultrafast endocytosis requires seconds (15). In the same neurons, VGLUT2-pH shows no effect of the β3B KO at any frequency in the absence of bafilomycin (Fig. 3B), and the β3B KO has a larger effect on VMAT2-pH in the absence than the presence of bafilomycin (Figs. 2A, 3A). Within the same cell population, loss of β3B thus has a selective effect on exocytosis of VMAT2^+^, not VGLUT2^+^ SVs. Inactivation of VPS41 specifically in dopamine neurons produces a similar, frequency-dependent defect in externalization of VMAT2 (Fig. 3C). The results indicate that AP-3 and VPS41 are required specifically for the exocytosis of monoamine SVs in response to high frequency stimulation. They also suggest that, in addition to VMAT2, AP-3 and VPS41 direct the trafficking to SVs of proteins required for exocytosis at high frequency. However, AP-3 does not simply target these proteins to a single, common pool of SVs, because if it did, loss of AP-3 should similarly impair the regulated exocytosis of VGLUT2^+^ vesicles. Since loss of AP-3 has minimal or no effect on VGLUT2, AP-3 must direct the formation of a novel SV population that both preferentially stores monoamine and responds specifically at high frequency. Consistent with a role for sorting to different SVs in the properties of release, loss of β3B does not affect Ca^++^ entry (*SI Appendix*, Fig. S5), and this as well as the lack of effect on VGLUT2^+^ SVs argues against effects of the KO on action potential invasion of the axon terminal. Although the trafficking of VGLUT2 does not depend on AP-3, other neurotransmitter transporters may rely on AP-3 for targeting to SVs, endowing their cognate transmitters with the capacity for phasic release. Indeed, the analysis of vesicular GABA transporter (VGAT)-pH shows an AP-3-dependent defect in release also specific to high frequency stimulation (*SI Appendix*, Fig. S6).

**Fig. 3.**
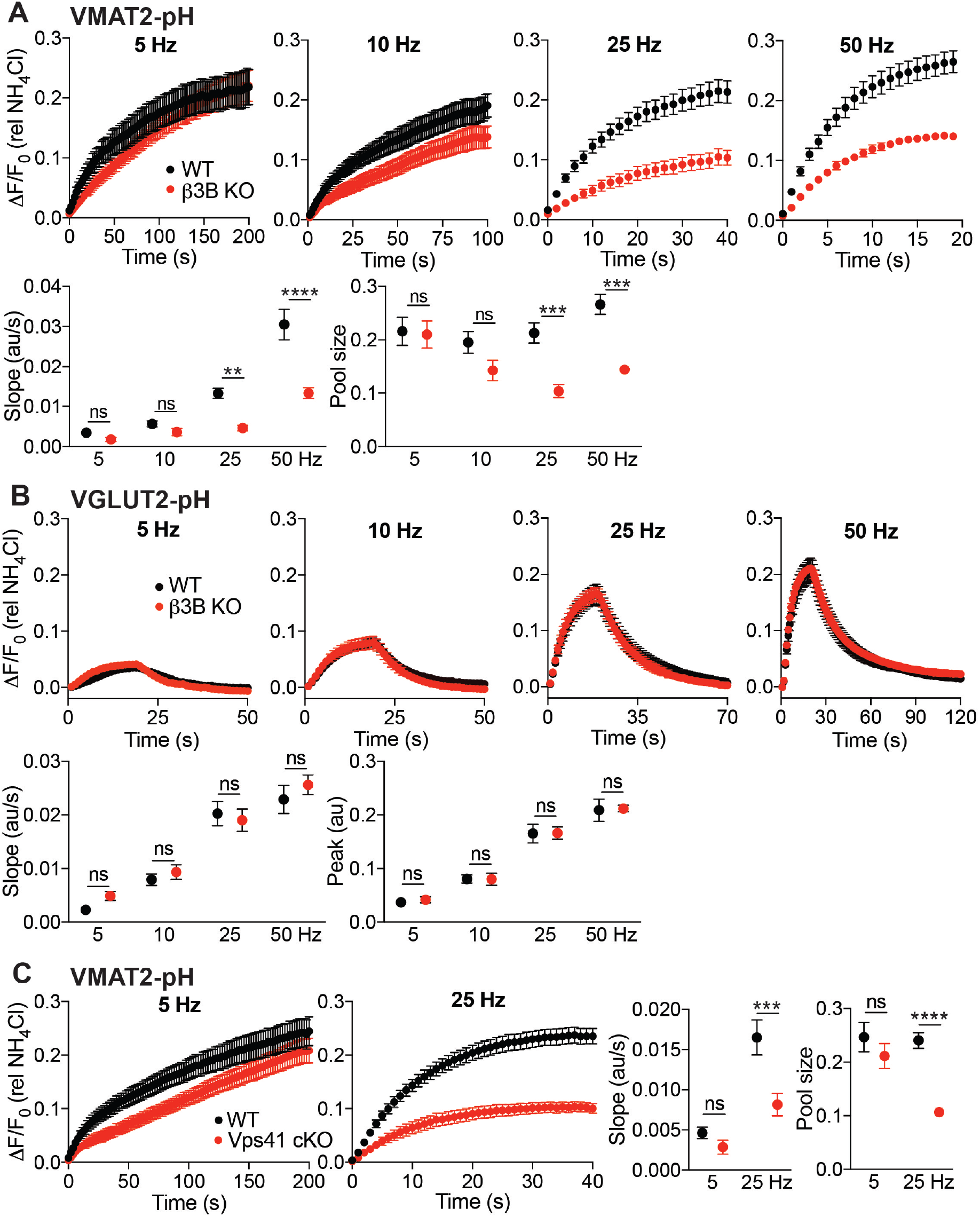
Loss of neural AP-3 preferentially impairs the response of VMAT2^+^ synaptic vesicles to high frequency stimulation. Midbrain dopamine neurons expressing VMAT2-pH were stimulated in the presence of 100 nM bafilomycin with 1000 action potentials at 35° C (A,C). Dopamine neurons expressing VGLUT2-pH were stimulated without bafilomycin for 20 s at 35° C (B). The fluorescence intensity was normalized to that in NH_4_Cl. (A) Cultures from WT and β3B KO mice were stimulated at different frequencies and the mean initial slope and pool size are shown below (n=9-10 coverslips for WT for β3B KO from 3 different cultures). (B) Cultures from WT and β3B KO mice were stimulated at different frequencies and the mean initial slope and peak are shown below (n=9 coverslips from 2 different cultures). (C) Cultures from WT and Vps41 cKO mice were stimulated at 5 and 25 Hz, the mean pool size and initial slope are shown to the right (n=9-10 coverslips for WT and Vps41 cKO from 3 different cultures). ***, p<0.001; ****, p<0.0001; ns, non-significant by two-way ANOVA with Sidak’s multiple comparisons test. Error bars indicate SEM.

### AP-3 Confers the Frequency Dependence of Striatal Dopamine Release

To determine whether the loss of neural AP-3 impairs the frequency dependence of dopamine release, we used cyclic voltammetry to monitor dopamine release in the dorsal striatum. In slices with multiple pre- and postsynaptic receptors blocked to prevent indirect effects on release, β3B KO mice show an ∼50% reduction relative to wild-type in peak dopamine release triggered with a single action potential (Fig. 4A), consistent with the observed reduction in striatal dopamine stores (Fig. 1A). We also find no change in the time constant of decay, indicating that the mutation does not impair the activity of DAT (Fig. 4A). At 5 Hz stimulation, the β3B KO shows a slightly greater reduction in peak dopamine that persists even after normalization for the reduced response to a single stimulus (Fig. 4B,D). At 25 Hz stimulation, the decrease after normalization to a single stimulus is substantially larger (Fig. 4C,D). At both frequencies, we used 25 action potentials, short trains closer to the burst firing of dopamine neurons and the role of neural AP-3 appears within one second, again suggesting relevance for the short bursts of dopamine neurons and arguing against a role restricted to SV regeneration. Thus, loss of AP-3 impairs dopamine release in two ways. First, it reduces evoked release by ∼50%, due to reduced dopamine stores. Second, loss of AP-3 further impairs the release of dopamine at high frequency due to its role in the biogenesis of a specific SV subpopulation that differs from canonical SVs in its low release probability. A simple reduction in the number of SVs would not affect the kinetics of release but we find that the residual release observed at high frequency in the β3B KO shows faster kinetics than WT (Fig. 4E), consistent with the selective loss of low release probability vesicles.

**Fig. 4.**
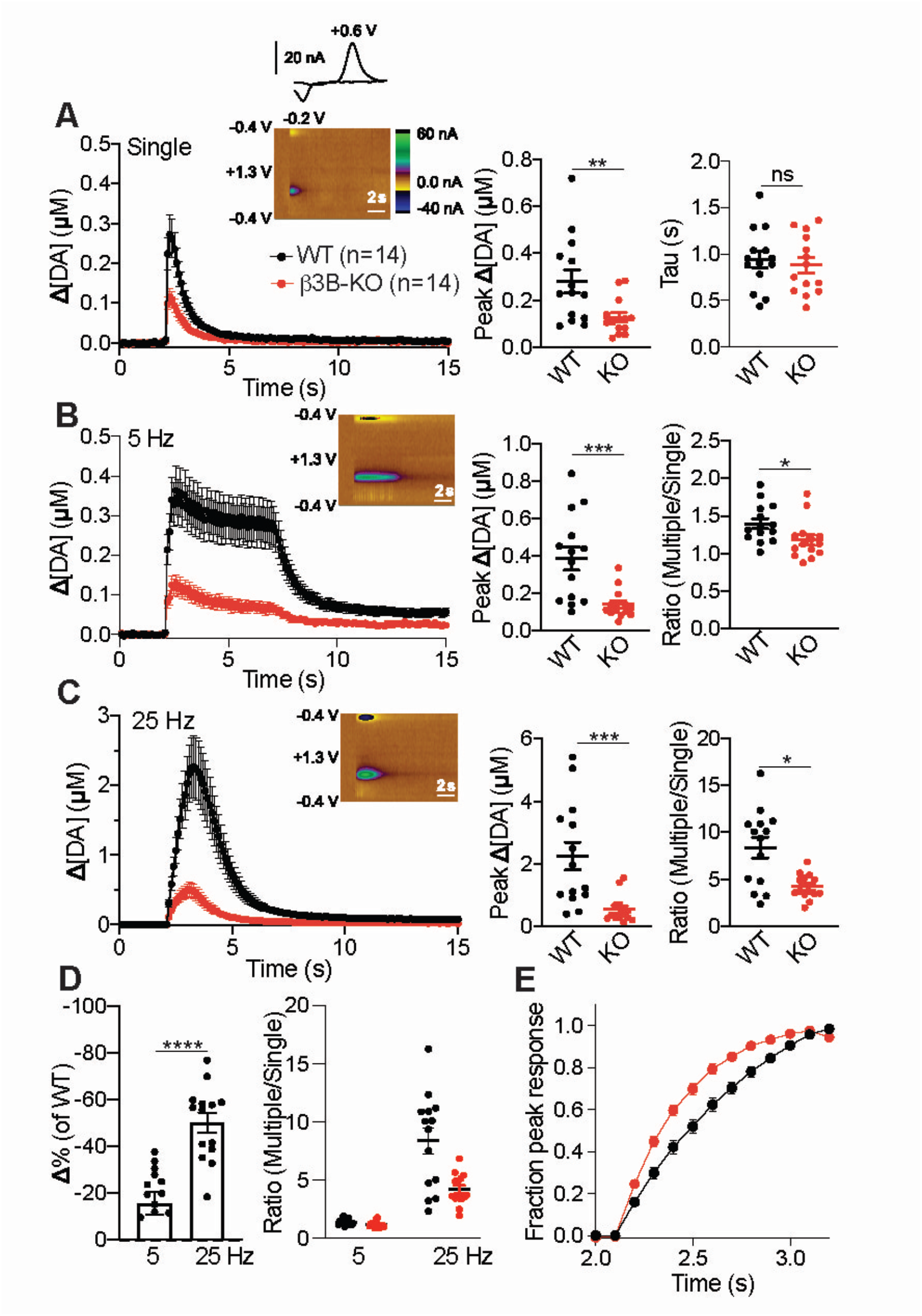
Neural AP-3 is required for striatal dopamine release in response to high frequency stimulation. (A) Time course of striatal dopamine release evoked by single stimuli and measured with FSCV. Inset: color plot (left) and characteristic dopamine voltammogram showing oxidation and reduction peaks at +0.6 V and −0.2 V, respectively. Scatterplot (middle) shows peak dopamine concentrations in β3B KO (n=14 from 4 animals) and WT littermates (n=14 from 4 animals). **, p<0.01, U=42, Mann-Whitney U-test. Scatterplot (right) shows the time constants of dopamine clearance (p=0.623 by t-test). ns, not significant. (B) Time course of dopamine release evoked by 25 stimuli delivered at 5 Hz. Scatterplots indicate peak dopamine concentrations (left) and ratio of 5 Hz/single stimulus (right). * p<0.05, U=44, Mann-Whitney U-test; ***, p<0.001, U=24, Mann-Whitney U-test. (C) Time course of dopamine release evoked by 25 stimuli delivered at 25 Hz. Peak dopamine concentrations (left) and ratio of 25 Hz/single (right). * p<0.05, U=46, Mann-Whitney U-test; ***, p<0.001, U=22, Mann-Whitney U-test. (D) Percent change of multiple/single stimulus ratios in β3B KO mice normalized to WT (left). ****, p<0.0001, t_(26)_=5.26 by independent samples t-test. Ratio of multiple (5 or 25 Hz) /single stimulus (right). ****, p<0.0001 by 2-way ANOVA. (E) Loss of neural AP-3 results in the acceleration of residual dopamine release at 25 Hz. The data in (C) are normalized to peak and fitted to a single exponential. The genotypes differ, p<0.0001. Error bars indicate SEM.

The role of AP-3 in the frequency dependence of dopamine release appears independent of its role in the polarity of dopamine storage, but it is possible that reduced dopamine storage affects the frequency dependence of release. To test this possibility, we used VMAT2 heterozygotes, which also show an ∼50% reduction in striatal dopamine stores (52, 53). Voltammetry in response to a single stimulus confirms reduced dopamine release by the VMAT2 heterozygote relative to wild-type (*SI Appendix*, Fig. S7A). Stimulation at 5 and 25 Hz produces a similar reduction in the VMAT2 heterozygote, with no additional impairment after normalization to the response to a single stimulus (*SI Appendix*, Fig. S7B,C). In further contrast to the β3B KO, the VMAT2 heterozygote shows no difference from WT in the kinetics of dopamine release at 25 Hz (*SI Appendix* Fig. S6D), consistent with a reduction in VMAT2 on all SVs. Thus, the reduced dopamine stores do not produce a frequency-dependent impairment of release like that observed in the β3B KO and VPS41 cKO, and the role of neural AP-3 and VPS41 in the frequency dependence of release is independent of their role in the axonal polarity of dopamine storage. Together with the differential role for AP-3 in monoamine and glutamate release (Fig. 2), the results suggest that AP-3 and VPS41 form a novel population of SVs that store monoamine and release preferentially in response to high frequency stimulation.

Considering the effects of AP-3 on the axodendritic polarity of VMAT2 trafficking, we also investigated somatodendritic dopamine release by voltammetry in the substantia nigra. The small amounts of dopamine released in the midbrain can make it difficult to distinguish from serotonin but the voltammetry does not show the peak characteristic of serotonin (54) (Fig. 5A). We find that despite the increased midbrain tissue dopamine, loss of β3B does not change the dopamine released by a single stimulus (Fig. 5A). In addition, the β3B KO does not alter the increase in somatodendritic dopamine release with 5 and 25 Hz stimulation (Fig. 5B,C). Thus, in contrast to the dorsolateral striatum, the frequency dependence of somatodendritic dopamine release does not depend on AP-3, suggesting that other proteins contribute to the formation of somatodendritic secretory vesicles. Since AP-3 confers the frequency dependence of release in the striatum, we then determined whether the AP-3-independent mechanism for release in the midbrain differs in sensitivity to stimulation frequency. Consistent with the difference in biogenesis, somatodendritic release in wild type mice appears less steeply dependent on the frequency of stimulation than striatal release (Fig. 5D).

**Fig. 5.**
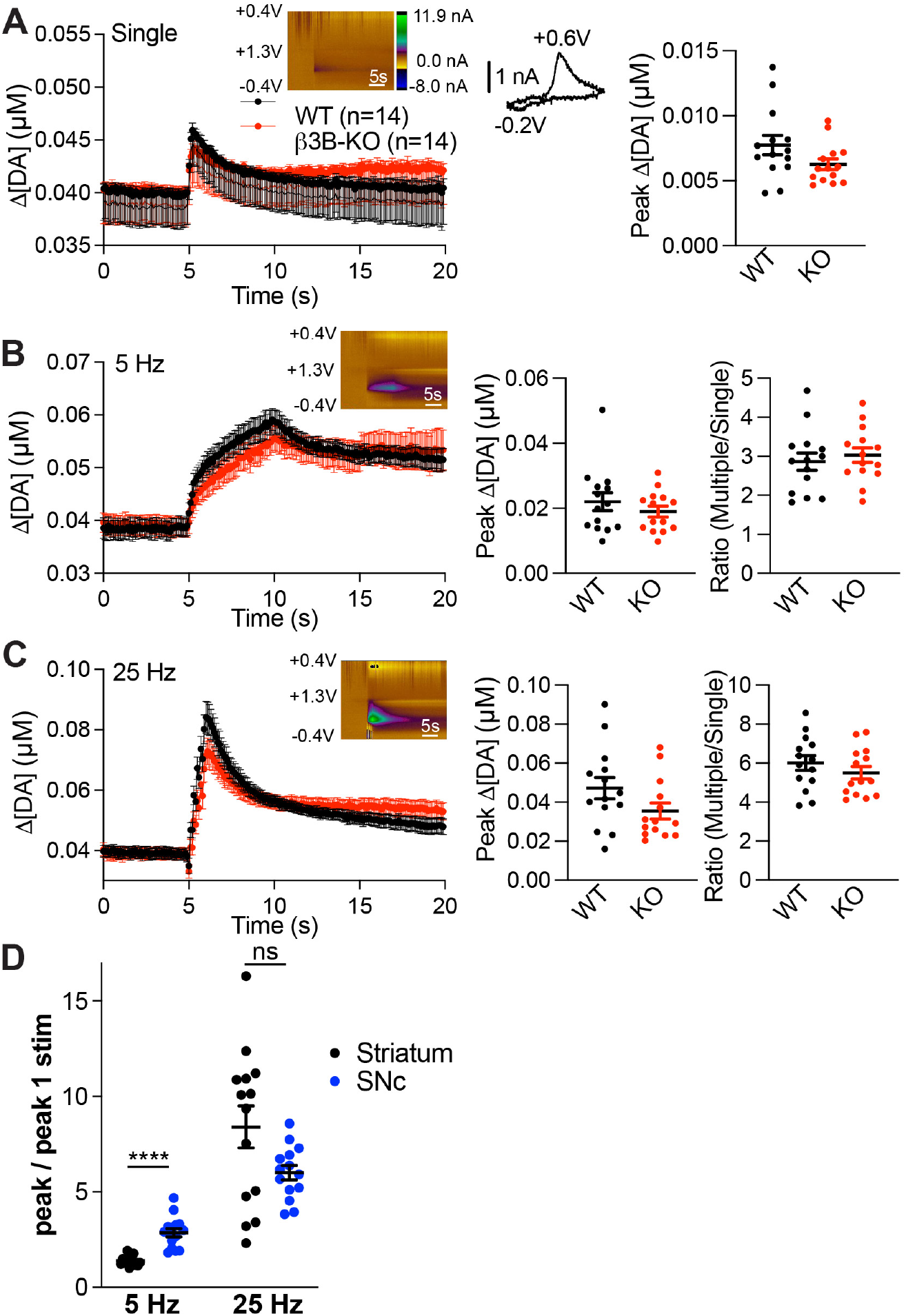
Loss of neural AP-3 does not alter substantia nigra dopamine release. (A) Time course of substantia nigra dopamine release evoked by single stimuli and measured with FSCV, as described in Fig. 4. Scatterplot shows peak dopamine concentrations in β3B KO (p=0.092). n=14 from 4 animals for β3B KO and WT littermates. (B) Time course of dopamine release evoked by 25 stimuli delivered at 5 Hz. Peak dopamine concentrations (left) (p=0.35) and ratio of 5 Hz/single stimulus (right) are shown as scatterplots (p=0.57). (C) Time course of dopamine release evoked by 25 stimuli delivered at 25 Hz. Peak dopamine concentrations (left) (p=0.0997) and ratio of 25 Hz/single (right) are shown as scatterplots (p=0.32). (D) Frequency dependence of dopamine release at 5 and 25 Hz in the striatum and substantia nigra, shown as scatterplots. ****, p<0.0001 by Mann-Whitney; p=0.15 at 25 Hz. Error bars indicate SEM.

### Conditional Inactivation of VPS41 in Dopamine Neurons Impairs Reinforcement Learning

The defect in striatal dopamine release with stimulation at high frequency suggested that loss of AP-3 may impair the learning dependent on dopamine released by burst firing (3). The reduction in dopamine release may also impair movement (55), but we find that the β3B KO does not reduce movement velocity, which increases slightly (*SI Appendix*, Fig. S8). To test a role in reinforcement learning, we used T-maze tasks in which learning was previously shown to require phasic dopamine release (56) and cKO mice with VPS41 selectively inactivated in dopamine neurons. In a task with the food reward consistently in one arm of the maze (Fig. 6A), VPS41 cKO mice exhibit a delay in learning relative to wild-type (Fig. 6B). In a second task with the reward linked to a visual cue but placed pseudorandomly (50:50) in either arm of the maze, VPS41 cKO mice also exhibit a substantial delay in learning (Fig. 6C). However, both AP-3 KO and VPS41 cKO mice show reduced striatal dopamine, raising the possibility that this rather than a change in the properties of release may account for the defect in learning. To test this possibility, we again took advantage of the VMAT2 heterozygotes, which show a reduction in dopamine levels similar to the β3B KO and the VPS41 cKO (52, 53) but without an effect on the frequency dependence of release (*SI Appendix*, Fig. S7). In contrast to the VPS41 cKO, VMAT2 heterozygotes show no learning defect in the uncued and cued T-maze tasks (Fig. 6D,E). Consistent with previous work using dopamine-deficient mice (57), a reduction of this magnitude in the amount of dopamine stored thus does not impair learning in the T-maze. The learning phenotype must therefore involve the defect in high frequency release of dopamine observed in AP-3 and VPS41 cKO mice. What accounts for the learning that remains in the VPS41 cKO? Global loss of β3B impairs learning to a similar extent as the Vps41 cKO in the uncued T-maze (Fig. 6F) but eliminates learning over the time course of training in the more difficult cued task (Fig. 6G). A role for AP-3 in the frequency-dependent release of other transmitters may thus account for the cued learning that remains after inactivation of this mechanism specifically in dopamine neurons.

**Fig. 6.**
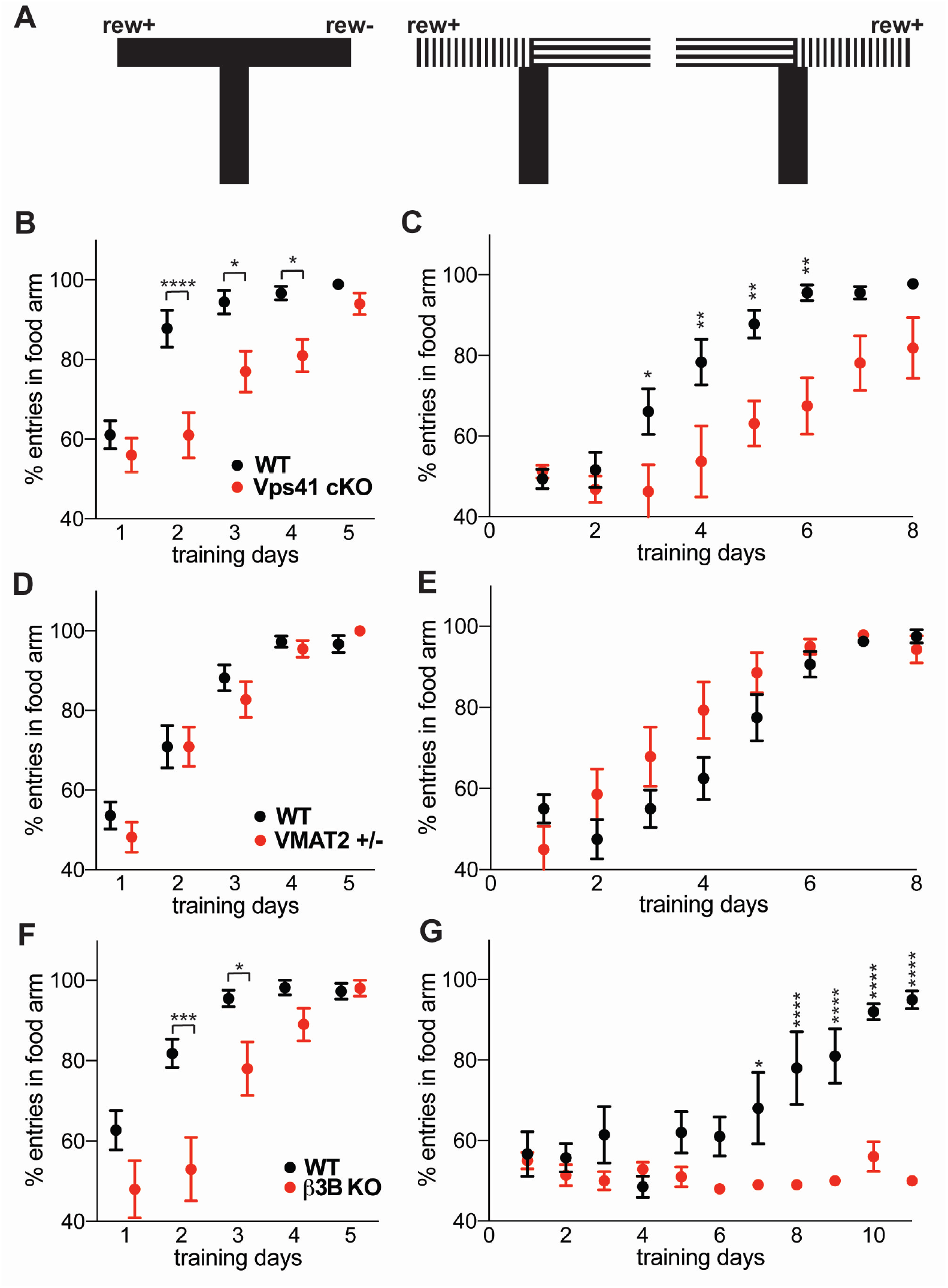
Inactivation of VPS41 in dopamine neurons impairs learning in the T-maze. (A) Schematic representation of the T-maze task. In the uncued T-maze task (left), food reward was always provided in the L arm. In the cued T-maze task (right), the reward was pseudorandomly (50:50) provided in either arm but always in association with vertical stripes. Each animal performed 10 trials per day in the uncued task and 20 trials per day in the cued, except for the first four days of testing in the β3B KO, where 14 trials were performed per day before increasing to 20. (B) Percent correct entries of WT (DATicre+/-) and VPS41 cKO mice into the food arm on each day of training in the uncued T-maze task (p=0.0331 for difference in genotype by 2-way ANOVA with Sidak’s *post hoc* multiple comparisons test). n=9 WT (DATicre+/-) and n=10 Vps41 cKO mice. (C) Percent correct entries by WT (DATicre+/-) and VPS41 cKO on each day of training in the uncued task (p=0.045 for genotype difference by 2-way ANOVA). n=9 WT (iDATcre+/-) and n=8 Vps41 cKO mice. (D) Learning by VMAT2 heterozygotes (+/-) in the uncued T-maze task (p=0.80 for genotype difference by 2-way ANOVA). n=11 WT and VMAT2 +/-mice. (E) Learning by VMAT2 heterozygotes (+/-) in the cued T-maze (p=0.07 by 2-way ANOVA). n=8 WT and n=7 VMAT2+/-mice. (F) Percent correct entries by β3B KO mice in the uncued T-maze task (p=0.032 for genotype difference by 2-way ANOVA). n=11 WT and 10 β3B KO mice. (G) Percent correct entries by β3B KO in the cued T-maze task (p<0.0001 for genotype difference by 2-way ANOVA). n=5 WT and 5 β3B KO mice. *, p<0.05; **, p<0.01; ***, p<0.001; ****, p<0.0001 for each time point by two-way ANOVA with Sidak’s adjustment for multiple comparisons. Error bars indicate SEM.

## Discussion

Previous work has implicated AP-3 in presynaptic membrane trafficking but with unclear effects on neurotransmission. We now find that AP-3 has two distinct roles in transmitter release. First, AP-3 promotes the axonal polarity of dopamine storage and release, and this requires the neural rather than the ubiquitous adaptor complex. Recent work in *C. elegans* has shown a requirement for AP-3 in the assembly of presynaptic release machinery (25). In mammals, the nerve terminal forms in the absence of AP-3 (32), but we now find that the β3B KO impairs axonal delivery of VMAT2. Since loss of AP-3 does not affect axonal targeting of several other SV transporters (27), the complex appears responsible for trafficking a subset of SV proteins to the axon terminal. Localization of neural AP-3 to the cell bodies of midbrain dopamine neurons presumably enables sorting of these proteins into carrier vesicles destined for the axon.

In the absence of AP-3, midbrain dopamine rises due to increased VMAT2 in the cell bodies and dendrites of midbrain dopamine neurons. A mechanism distinct from AP-3 thus targets VMAT2 to the somatodendritic domain, and previous work has suggested a role for the related AP-1 complex in somatodendritic targeting (25, 58-60). Despite the increased stores, the β3B KO does not increase somatodendritic dopamine release, suggesting that in the absence of neural AP-3, VMAT2 redistributes to membranes capable of dopamine storage but not release.

Second, AP-3 confers the frequency dependence of dopamine release. The analysis of glutamate corelease by dopamine neurons has shown that AP-3 produces a subpopulation of SVs that store and release dopamine, with little effect on glutamate (13). We now find that loss of AP-3 preferentially impairs the exocytosis of VMAT2^+^ SVs and dopamine release in response to stimulation at high frequency, with relatively little effect at low frequency. The selectivity for VMAT2 rather than VGLUT2 argues against a role for AP-3 in a process required for externalization of both reporters such as spike propagation or energetics and in favor of a more specific role for AP-3 in the generation of SVs that respond at high frequency and contain little VGLUT2. In addition, the results indicate that VMAT2 targets to both the AP-3-derived subpopulation of low release probability SVs that respond to high frequency stimulation and the AP-3-independent subpopulation that responds at low frequency. The role of AP-3 in SV biogenesis also requires the neural complex, and, consistent with this specificity, the axon contains only neural AP-3, in contrast to the cell body and dendrites which also contain the ubiquitous complex.

Neural AP-3 presumably assembles the membrane proteins that endow SVs with low release probability. VAMP7 and the zinc transporter ZnT3 depend on AP-3 for trafficking to SVs (28, 32, 61), suggesting that they may be responsible. The zinc coreleased with glutamate does not appear to affect transmitter release (33). However, loss of VAMP7 increases neurite extension (62), suggesting an inhibitory role in membrane insertion that might also contribute to the low probability of transmitter release. Other proteins on and off SVs presumably contribute to the release properties of AP-3-derived SVs.

The role of neural AP-3 in SV formation from endosomes and the requirement for AP-3 in release at high frequency suggest that AP-3 produces a distinct subpopulation of SVs with low release probability. However, mice lacking AP-3 also show a reduced number of SVs (27), raising the possibility that SV depletion simply limits release more at high frequency. The faster kinetics of residual release at high frequency in the β3B KO argues against this possibility. A simple reduction in the number of all SVs would not change the kinetics of release and the unchanged kinetics of the VMAT2 heterozygote serve as control to demonstrate this. Loss of β3B thus eliminates a specific subset of SVs that responds with low release probability.

Loss of AP-3 reduces the pool of recycling SVs labeled with VMAT2-pH but also affects the initial rate of response, well before exhaustion of the recycling pool. In addition, the imaging of VMAT2-pH is normalized to the total pool of presynaptic reporter revealed in NH_4_Cl. The defect in release observed in the absence of AP-3 thus occurs independent of a reduction in the total pool of VMAT2-pH. Previous work has also shown an important role for the dopamine transporter DAT in the summation of dopamine release at high frequency stimulation (63). Increased DAT activity might therefore impair dopamine release specifically at high frequency, but we observe no change in dopamine clearance to indicate a change in DAT function. In addition, the ∼50% reduction in dopamine release observed at low frequency in the absence of β3B could influence the activation of presynaptic receptors to produce additional effects on release at high frequency, but we have used multiple receptor antagonists to prevent these effects. Further, the VMAT2 heterozygote, with a similar reduction in dopamine release as the β3B KO, shows no additional defect in dopamine release at high frequency. Thus, the role of neural AP-3 in the frequency dependence of dopamine release is independent of its role in the absolute amount of dopamine released. In contrast to the striatum, loss of β3B does not affect the frequency dependence of dopamine release in the midbrain. Thus, AP-3 does not appear to be involved in the formation of somatodendritic secretory vesicles.

VPS41 appears to serve as the coat protein for AP-3 in the formation of dense core vesicles (48, 64). We now find that VPS41 also contributes to the role of AP-3 in axonal polarity and SV biogenesis. Loss of VPS41 produces the same depletion of striatal dopamine as loss of β3B and preferentially impairs the response to stimulation at high frequency. Thus, VPS41 cooperates with neural AP-3 in multiple trafficking processes that contribute to neurotransmitter release.

The effect of AP-3 and VPS41 on SV exocytosis as well as recycling contrasts with the apparently more restricted role of AP-2 and clathrin in SV formation from the plasma membrane or by ultrafast endocytosis (15, 16 Kononenko, 2014 #6676). Loss of these factors has not been found to affect initial release (17, 65) as shown here for AP-3 and VPS41, suggesting that additional factors contribute to the composition and properties of canonical SVs made independent of AP-3.

The importance of AP-3 and VPS41 for dopamine release at high frequency suggested a role for the two proteins in reinforcement learning. To test this, we took advantage of a conditional KO, inactivating VPS41 specifically in dopamine neurons. The cKO mice show substantial delay in uncued as well as cued learning in the T-maze. Nonetheless, they eventually acquire the tasks, indicating preservation of basic behaviors such as ambulation. However, loss of either AP-3 or VPS41 reduce striatal dopamine content and this rather than a defect in the frequency dependence of release may account for the delay in learning. To distinguish between these possibilities, we took advantage of the VMAT2 heterozygotes, which also exhibit reduced striatal dopamine stores but no specific defect in release at high frequency. The VMAT2 heterozygote shows no defect in learning, demonstrating that the learning impairment observed with loss of VPS41 reflects the alteration in frequency dependence of release, not the reduction in dopamine stores. However, it may be difficult to extrapolate from the defect in release at high frequency observed *in vitro* directly to behavior. In striatal slices, we silenced essentially all receptors (such as cholinergic) that provide feedback onto dopamine release. *In vivo*, these signals will influence dopamine release and contribute to behavior.

The results have important implications for disorders of dopamine signaling. In Parkinson’s disease, the striatal content of dopamine is reduced. However, the results indicate that this alone would not disrupt reinforcement learning. Consistent with this, previous work has shown that a dramatic reduction in dopamine levels due to almost complete inactivation of VMAT2 does not produce parkinsonism until late in life (66). Thus, the amount of dopamine released may be less important for motor control and reinforcement learning than the appropriate frequency dependence of release. Indeed, homeostatic changes in receptor sensitivity and dopamine clearance would be expected to compensate for changes in the amount of dopamine released but not for an alteration in the frequency dependence of release.

The dopamine release that persists in the absence of AP-3 reflects the trafficking of VMAT2 to SVs not produced by AP-3 that respond to low frequency stimulation. We hypothesize that this SV population contributes to tonic dopamine release. Since VGLUT2 shows relatively little dependence on AP-3, it may target to the same AP-3-independent population of SVs, enabling the costorage of glutamate and dopamine. Indeed, previous work has demonstrated a role for VGLUT2 in the storage of dopamine by glutamate-coreleasing neurons in the ventral tegmental area, and this synergy requires expression of both transporters on the same SVs (37, 67). Further, the AP-3-dependence of VGAT and VMAT2 but not VGLUT2 suggests that neurotransmitter transporters slot onto AP-3-derived SVs in different proportions, contributing to different modes of release, and that the role of AP-3-derived SVs in phasic release extends beyond monoamines.

In summary, the results show how a protein originally implicated in endolysosomal trafficking in yeast has evolved to confer the frequency dependence of dopamine release and enable its role in reinforcement learning.

## Materials & methods

### Animals

Mice of both sexes were used. They received unlimited food and water under 12h light/12h dark cycles. Vps41^fl/fl^ mice (68) were crossed with DAT-ires-cre/DAT-ires-cre (DATicre; JAX stock 006660) and then with Vps41^fl/wt^;DATicre/-to generate Vps41^fl/fl^;DATicre/- and littermate controls (Vps41^fl/fl^). Heterozygous β3B mice (JAX stock 005684) were crossed with HA-VMAT2 BAC transgenic mice (31) to produce β3B KO/HA-VMAT2 animals. Wild-type (WT), β3B knockout (KO), HA-VMAT2, β3B KO;HA-VMAT2, Vps41^fl/fl^, Vps41^fl/fl^;Daticre^+/-^ mice that were 2-3 months old were used for anatomy and catecholamine measurements. β3B KO mice and WT littermates (3-6 months old) were used for the slice recording. β3B KO mice and WT littermates, Vps41^fl/fl^;Daticre^+/-^ and DATicre^+/-^, VMAT2 +/- and WT littermate (2-6 months old) were used for T-maze experiments. All animal experiments were approved by the University of California San Francisco Institutional Animal Care and Use Committee and performed as required by the National Institutes of Health Guide for Care and Use of Laboratory Animals. For most of the voltammetry experiments, mice were transferred to the University of Colorado, where the procedures were approved by the University of Colorado School of Medicine and conducted in accordance with the protocols approved by the National Institutes of Health Guide for Care and Use of Laboratory Animals.

### DNA construct

VGAT-pHluorin was amplified from pCAGGS VGAT-pHluorin (Addgene plasmid # 78578) using XbaI and NheI sites, and cloned into the lentivirus vector pFUGW. The following primers were used for the PCR amplification: forward primer: 5’ GCG TCT AGA CCC TTG CTG CCT TGA CGC GCG CC; reverse primer: 5’ CGC GCT AGC TAG GCG TAG TCC GGC ACG TCG TAC. The final construct was verified by DNA sequencing.

### Lentivirus production

HEK293T cells were transfected with third-generation lentiviral vector pFUGW encoding the gene of interest + accessory plasmids pREV, pVSVG and pPRE), using a Fugene transfection reagent (Promega; Cat# PRE2311). The medium was replaced the next day and lentivirus collected 36 h later, with cell debris removed by centrifugation at 1000g. The virus supernatant was either used immediately or stored at −80° C for future use.

### Primary culture

#### Hippocampal neurons

Hippocampi were dissected at postnatal day 0 from WT, β3A KO, β3B KO and *mocha* mice in Hank’s balanced salt solution (HBSS) containing 10 mM HEPES and 20 mM glucose. Hippocampi were dissociated with papain (20 units/mL) in papain digestion buffer (HBSS containing 20 mM CaCl_2_, 5 mM EDTA, 0.2 mg/mL L-Cysteine, 10 mM HEPES, pH 7.4), washed three times with HBSS containing 10 mM HEPES and 20 mM glucose, triturated, and then plated onto poly-L-lysine-coated coverslips in Minimal Essential Medium containing 1x B27 (GIBCO, 17504-044), 2 mM glutamax (GIBCO, 35050-061), 5% FBS (HyClone, defined), 21 mM glucose (Sigma, G8769) at a density of 350 cells/mm^2^. After day 1 *in vitro* (DIV1), 3/4 of the medium was replaced with Neurobasal medium (GIBCO, 21103-049) containing B27 and glutamax. Cells were infected at DIV4-5 with lentiviruses encoding VGLUT2-pH, VMAT2-pH or VGAT-pH. Glial proliferation was inhibited with 4 μM Ara-C added on DIV6-7.

#### Dopamine neurons

The midbrain (including ventral tegmental area and substantia nigra) was dissected from P0 pups in HBSS containing 10 mM HEPES and 20 mM glucose, digested for 15 minutes with papain (20 units/mL) in papain digestion buffer, washed three times with HBSS containing 10 mM HEPES and 20 mM glucose, triturated and plated at 1000 cells/mm^2^ in medium containing 60% Neurobasal medium, 30% Basal Eagle Medium, 10% fetal bovine serum (HyClone, defined), 1x B27, 2 mM GlutaMAX, 10 ng/mL glial cell line-derived neurotrophic factor (Gibco PHC7045) and 1x penicillin/streptomycin onto an astrocyte monolayer itself previously plated onto poly-L-lysine and laminin-coated glass coverslips. The dopamine neurons were infected with lentiviruses encoding VMAT2-pH at DIV 2-4.

### Measurement of catecholamine levels

Mice were euthanized by inhalation of CO_2_, decapitated and the brain removed. The brains were placed into a rodent brain matrix (RBM-2000C, Protech International Inc.), two 1 mm coronal slices cut, cortical tissue removed, and striatal tissue collected in cold HBSS containing 10 mM HEPES and 20 mM Glucose. The midbrain containing the ventral tegmental area and substantia nigra was dissected from the same brain. All the collected tissue was flash-frozen in liquid nitrogen and immediately transferred to −80° C. Tissue catecholamine levels were measured by HPLC with coupled electrochemical detection at the Vanderbilt Neurochemistry Core.

### [^3^H]-Dihydrotetrabenazine binding

The midbrain and striatum were isolated using rodent brain matrix as described above, collected into cold SHT buffer (320 mM sucrose, 10 mM HEPES/Tris pH 7.4) with 0.5 mM EDTA and Complete Protease Inhibitor Cocktail (Roche). As described previously(13), tissue from midbrain and striatum was disrupted with 12 strokes of a Dounce homogenizer at 500 rpm in cold SHT buffer and sonicated for 30 s before sedimentation of the debris at 2000*g* for 2 min. The protein content of the supernatant was determined by BCA and 50 μg membrane protein incubated in SHT buffer with 10 nM (+)-a-dihydrotetrabenazine [9-O-methyl-^3^H] (ARC; 80 Ci/mmol) for 30 minutes at 30° C. The reaction was stopped by filtration through a Supor 200 0.2 μm filter (PALL) and washed 3 times in ice-cold SHT buffer with 20 mM tetrabenazine (Fluka). Binding was performed in triplicate for each sample and nonspecific binding determined with 10 μM non-radioactive tetrabenazine added to the binding assay. Specific binding was normalized to the amount of membrane protein added to the reaction.

### Immunohistochemistry

Mice were perfused with PBS and then with 4% PFA in PBS. The dissected brains were fixed overnight at 4° C in 4% PFA/ PBS, cryoprotected in 30% sucrose in PBS overnight at 4° C and 35 µm sections cut using a Leica CM3050 S cryostat. Cultured neurons were fixed for 10 minutes at room temperature in PBS containing 4% PFA and 4% sucrose. Brain sections as well as cultured neurons was washed three times in PBS, blocked and permeabilized at room temperature in PBS containing 4% normal goat serum and 0.2% Triton X-100. For brain slices, the primary and secondary antibodies were diluted in PBS containing 1% normal goat serum and 0.2% Triton X-100. For cultured neurons, the antibodies were diluted in PBS containing 1% normal goat serum. Both brain slices and cultured neurons were incubated in primary antibody overnight at 4° C, washed three times, incubated with secondary antibodies at room temperature for 1 hour in the dark, and then washed three times. Chicken MAP2 antibody was used at 1:2000, mouse and rabbit anti-TH at 1:1000, and all other primary antibodies at a dilution of 1:500. All secondary antibodies were used at 1:1000. After staining, the brain sections and cultured neurons were mounted in Fluoromount-G (SouthernBiotech), and imaged using the Nikon Ti Microscope equipped with CSU-W1 spinning disk confocal, Plan Apo VC 100x/1.4 oil or 60x/1.4 oil objective and Andor Zyla sCMOS camera. A Nikon 6D conventional wide-field microscope, Plan Apo 20x/0.75 air objective and DS-Qi2 monochrome camera were used to stitch multi-channel fluorescence images of whole brain sections. Fluorescence images were analyzed using ImageJ (National Institutes of Health, Bethesda, MD). In brain slices, dendrites and representative sections of the axon were traced by one-pixel lines, and fluorescence intensity of VMAT2-HA and TH per µm process length from 20 processes used to determine the mean intensity per field. Values are represented as the mean, and error bars represent SEM. In cultured neurons, axons and dendrites were also traced by one-pixel lines and the mean intensity per µm process length determined in either three segments of the same axon or in three separate dendrites of the same neuron.

### Live imaging

Hippocampal and dopamine neurons were imaged at, respectively, DIV14-16 and DIV13-15. To label dopamine neurons, midbrain cultures from wild-type, β3B KO, Vps41^fl/fl^ and Vps41^fl/fl^;Dat-ires-cre^+/-^ mice were incubated for 5 min in Tyrode’s buffer (119 mM NaCl, 25 mM HEPES, 2 mM CaCl_2_, 2 mM MgCl_2_, 2.5 mM KCl, and 30 mM glucose at pH 7.4) containing fluorescent DAT ligand JHC 1-64 (30 nM)(13). The cultures were then rinsed in Tyrode’s buffer and both hippocampal and midbrain cultures mounted in the perfusion and stimulation chamber of a TE300 inverted epifluorescence microscope. The fluorescence of pHluorin was monitored using 470/40 nm excitation and 525/70 nm emission bandpass filters. The fluorescence of JHC1-64 was monitored with 545/25 nm excitation and 605/70 nm emission bandpass filters. After identifying dopamine neurons in the red channel using JHC 1-64 fluorescence, we switched to the pHluorin channel. In most experiments, we acquired images at 1 Hz. Stimulation was done by evoking action potentials (APs) with 1 ms bipolar current pulses through platinum-iridium electrodes at 5-10 V/cm. Cells were continuously perfused in Tyrode’s buffer with 10 μM 6-cyano-7-nitroquinoxaline-2,3-dione (CNQX) and 10 μM 3-(2-carboxypiperazin-4-yl) propyl-1-phosphonic acid (APV). Imaging was done at room temperature, except for the frequency dependence experiments in dopamine neurons where the imaging was done at 35° C. The fluorescence was normalized to total bouton pHluorin, which was determined by perfusing with Tyrode’s buffer containing 50 mM NH_4_Cl (and NaCl reduced to 69 mM). To eliminate the contribution of endocytosis, cells were continuously perfused with Tyrode’s solution containing 100 nM Bafilomycin A1 (Cayman, 11038). Analysis of the images was performed using ImageJ software (National Institutes of Health). Synaptic boutons were manually selected from the images in NH_4_Cl. Background fluorescence was subtracted, and the mean fluorescence intensity of the synaptic boutons normalized to pre-stimulation baseline and to unquenched (total) pHluorin fluorescence in NH_4_Cl. The data points were fitted using GraphPad Prism. To calculate initial slope, the fluorescence trace for the initial 4 seconds was fitted to a line and slope was calculated from this fit. Pool size was determined as fluorescence amplitude observed after the stimulation. In all cases, the fluorescence was normalized to that observed in NH_4_Cl.

### Fast scan cyclic voltammetry

Coronal brain slices (240 µm) containing the striatum were prepared from β3B KO mice and WT littermates (3-6 months old). For striatal sections, mice were deeply anesthetized with isoflurane and trans-cardially perfused with cold cutting solution containing (in mM): 75 NaCl, 50 sucrose, 6 MgCl_2_, 2.5 KCl, 1.2 NaH_2_PO_4_, 0.1 CaCl_2_, 25 NaHCO_3_, and 2.5 D-glucose (bubbled with 5% CO_2_ in O_2_). The brain was quickly removed and sectioned using a vibratome (Leica VT1200S) before slices were transferred to an incubation chamber filled with artificial cerebrospinal fluid (ACSF) containing (in mM): 126 NaCl, 2.5 KCl, 1.2 MgCl_2_, 1.2 NaH_2_PO_4_, 2.5 CaCl_2_, 21.4 NaHCO_3_, 11.1 D-glucose and 10 µM MK-801 (34° C, bubbled with 5% CO_2_ in O_2_, pH 7.4). After equilibration (at least 45 min), slices were individually transferred to a recording chamber. NMDA, AMPA, GABA_A_, GABA_B_, nicotinic, muscarinic, D2 and D1 receptors were blocked using 10 µM MK801, 10 µM DNQX, 10 µM picrotoxin, 0.3 µM CGP55845, 1 µM DHBE, 0.3 µM scopolamine, 0.5 µM sulpiride and 10 µM SCH23390, respectively. For midbrain sections, the mice were anesthetized with ketamine/xylazine and perfused with (in mM): 10 NaCl, 180 sucrose, 7 MgCl_2_, 2.5 KCl, 1.25 NaH_2_PO_4_, 0.5 CaCl_2_, 25 NaHCO_3_, and 10 D-glucose (bubbled with 5% CO_2_ in O_2_). Horizontal sections (240 µM) through the midbrain were prepared with a Leica VT1000S before transfer to modified ACSF containing (in mM): 125 NaCl, 2.5 KCl, 1.0 MgCl_2_, 1.25 NaH_2_PO_4_, 2.0 CaCl_2_, 25.0 NaHCO_3_, 10 D-glucose) and bubbled with 5% CO_2_ in O_2_, pH 7.4 at 34° C for 30 min to recover, followed by room temperature. The inhibitors used were the same as those used for striatal recordings except that the midbrain recordings did not include MK-801. All recordings were made at 31° ±1° C.

Electrically stimulated dopamine release was measured by fast-scan cyclic voltammetry (FSCV) using a carbon fiber microelectrode (CFM; 34-700, Goodfellow) with exposed diameter of 7 µm and length 50-90 µm. Microelectrodes were positioned in the dorsal striatum or substantia nigra pars compacta and scanned across a triangular waveform (−0.4 to +1.3 V, 400 V/s) at 10 Hz. Dopamine release was evoked using a monopolar glass stimulating electrode. In striatal sections, single (30 µA, 0.5 ms) or multiple (25x) pulses were delivered at 5 or 25 Hz. In the substantia nigra, stimuli were delivered at 250 µA for 1 ms. Release evoked by single pulse stimulation was averaged from 3 sweeps after stabilization (inter-sweep interval, ISI = 300 s). Release evoked by trains of stimuli was measured from a single sweep (ISI = 300 s). Non-faradaic dopamine signals were calculated by subtracting background currents averaged from the first 10 scans from each sweep. The time-course of release was measured at the oxidation peak of dopamine (+0.6 V). CFMs were calibrated after each experiment in a beaker containing Tris (10 mM)-buffered ACSF (pH 7.4) on a magnetic stirrer. Aliquots of dopamine were sequentially added and a second order polynomial was used to interpolate dopamine concentrations (µM) from peak oxidation currents (nA).

### Open Field Behavior

Adult mice (β3B KO or WT littermates) were habituated to the open field chamber (a 25 cm diameter clear acrylic cylinder) for at least 30 minutes/day over 2 sessions. Subsequent open field sessions were 30 minutes in duration. Mouse movement was monitored with an overhead camera and commercial video tracking software (Noldus Ethovision). Average movement velocity was calculated as the total distance traveled, divided by 30 minutes. The experimenter was blinded to the genotype of the mice during testing.

### T-maze tasks

As described previously (69), the mice were handled for one week and calorie-restricted to ∼85% of their original body weight, measuring their weight daily. To familiarize them with the food reward, they were given 10 reward pellets in their home cage. To familiarize them with the T-maze and the associated food reward, each mouse was allowed to explore the T-maze freely for 10 min with 5 rewards in each arm, and this was repeated over two days. In the simple, uncued task, food was always provided in the left arm of the T-maze. For testing, mice were placed in the T-maze start area for 60 sec. The sliding door was then removed and mice were given 60 sec to choose between the left arm that contained food (reward arm) and the right arm that did not (non-reward arm). After the choice was made, a door was placed in front of the arm entered and the mice had 60 sec to consume the food (reward) pellet, which they all did. If a mouse failed to make a choice within 60 sec, it was forced on an alternating basis into one of the arms, and held there for 60 sec. In the uncued task, ten trials were performed on each testing day. For the cued T-maze task, a mold was placed outside of the T-maze arms, with vertical stripes indicating the food arm and horizontal stripes the non-food arm. The task was performed as above, but the striped visual cue (always associated with food reward) was assigned pseudo-randomly to left and right arms, maintaining a 50:50 ratio overall. In this cued task, 20 trials were performed each testing day except in the experiment involving β3B knockouts where 14 trials were performed for each for the initial four days, then the number of trials increased to 20 per day.

### Statistical methods

Pairwise comparisons were evaluated by Mann-Whitney test. Multiple comparisons were evaluated by either one-way ANOVA with *post hoc* Bonferroni’s multiple comparisons test or two-way ANOVA with *post hoc* Sidak’s multiple comparisons test. Data are presented as mean ± SEM, and statistical significance as: * p<0.05, ** p<0.01, *** p<0.001, and **** p<0.0001. All of the analysis was performed using GraphPad Prism (GraphPad Software, La Jolla, CA).

## Acknowledgments

We thank the members of the Edwards lab for discussion and suggestions, J. Berke, L. Zweifel and R. Malik for help with the behavioral studies.

## Funding

National Institute of Neurological Disease and Stroke R01NS103938 (RHE)

Aligning Science Across Parkinson’s Disease ASAP-020529 (RHE, CPF, AN)

National Institutes of Health grant R01-DA35821 (CPF), R01-NS95809 (CPF)

National Institute on Drug Abuse F32-DA051135 (AY)

American Heart Association 15POST25750061 (SJ)

Brain & Behavior Research Foundation (JWM, KS)

## Data sharing plans

All study data are included in the article and/or SI Appendix. Data shall be made available upon reasonable request to R.H. Edwards, Departments of Physiology and Neurology, UCSF School of Medicine, 600 16^th^ St., GH-N272B, San Francisco, CA 94143.

**Competing interests:** RHE is a consultant for Nine Square Therapeutics. The authors declare that they have no other competing interests.

**Correspondence and requests for materials** should be sent to R.H. Edwards, Departments of Physiology and Neurology, UCSF School of Medicine, 600 16^th^ St., GH-N272B, San Francisco, CA. 94143.

**Data sharing plans:** All study data are included in the article and/or SI Appendix. Data shall be made available upon reasonable request from R.H. Edwards, Departments of Physiology and Neurology, UCSF School of Medicine, 600 16^th^ St., GH-N272B, San Francisco, CA. 94143.

## Notes

### Competing Interest Statement

The authors have declared no competing interest.

