## Supplementary material for "Adaptor Protein-3 Produces Synaptic Vesicles that Release Phasic Dopamine": SI

Classification: Biological Sciences, Neuroscience

Keywords: phasic dopamine; reinforcement learning; adaptor protein-3 (AP-3), VPS41, vesicular monoamine transporter 2 (VMAT2); synaptic vesicles

### Supplemental Figures

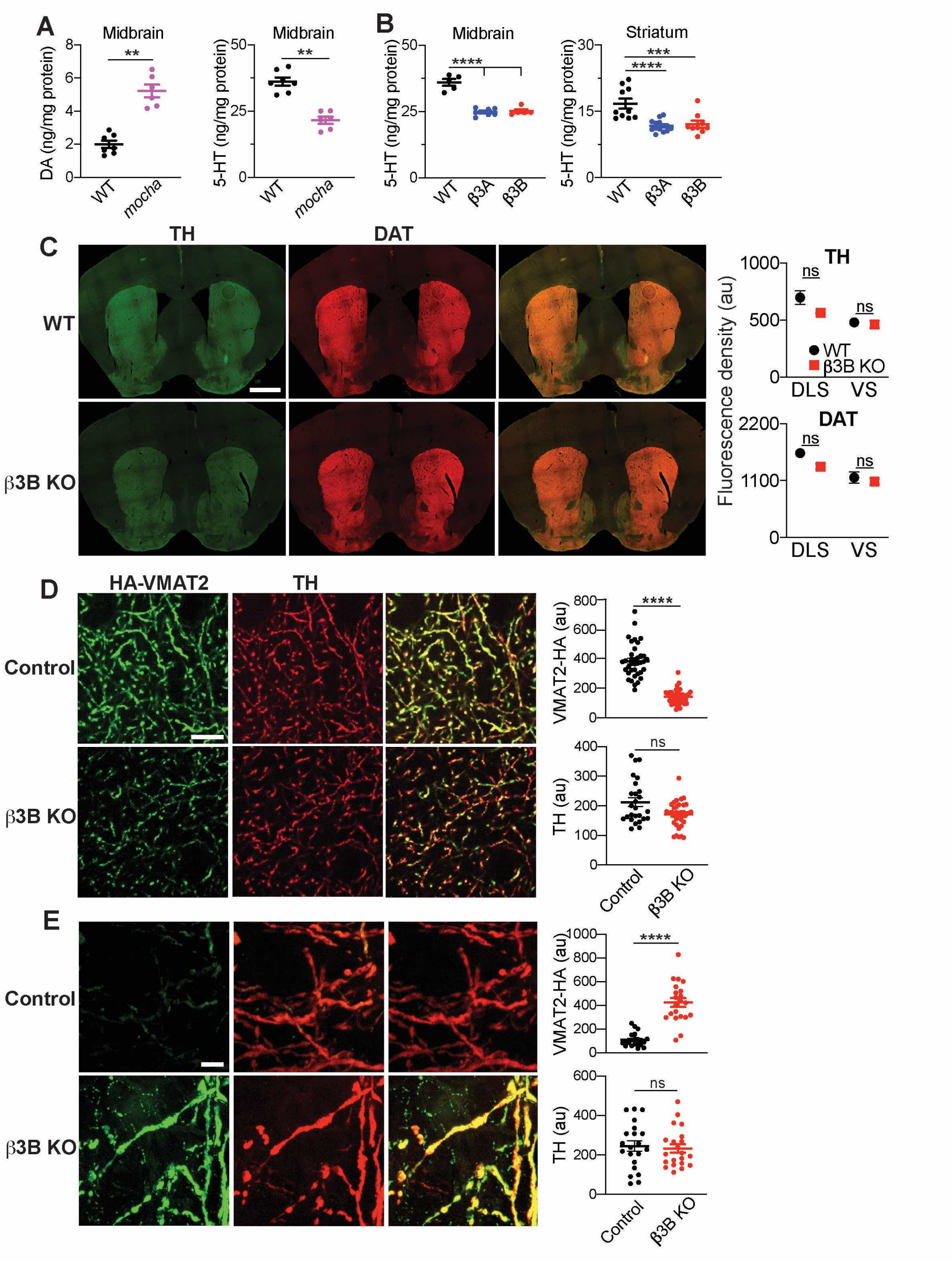

**Fig. S1. Loss of AP3 reduces DA and serotonin (5-HT) storage *in vivo*. The neural isoform of AP-3 is required for the polarity of VMAT2 localization *in vivo*.**

(A) The *mocha* mutation increases midbrain DA and reduces midbrain 5-HT. **, p=0.0012, by Mann-Whitney test (n=7 mice for WT and 6 for mocha). (B) Loss of either the ubiquitous isoform β3A or neuronal isoform β3B reduces 5-HT in midbrain and striatum. ***, p<0.001; ****, p<0.0001 by one-way ANOVA with Bonferroni's *post hoc* test (n=5 mice for WT, 7 for β3A KO and 5 for β3B KO in midbrain; n=10 for WT, 14 for β3A KO and 9 for β3B KO in the striatum). (C) Striatal slices from WT and β3B KO transgenic mice were immunostained for TH (green) and DAT (red). The mean fluorescence density per $\mu m^2$ area of individual fields is shown in the scatter plots. DLS, dorsolateral striatum, VS, ventral striatum. ns, not significant by two-way ANOVA with Sidak's multiple comparisons test (n=6 fields from three mice for each condition). Scale bar, 1 mm.

(D) Striatal slices from WT and β3B KO HA-VMAT2 BAC transgenic mice were immunostained for HA (green) and TH (red). Scatterplots (right) show mean fluorescence per μm process length for different fields. ****, p<0.0001; ns, not significant by Mann-Whitney test (n=37 fields from 3 WT mice and 42 fields from 3 β3B KO mice, both expressing the HA-VMAT2 BAC transgene; and n=26 fields from 3 WT mice, 34 fields from 3 β3B KO mice for TH). Scale bars, 5 μm. (E) Midbrain slices from WT/β3B KO HA-VMAT2 BAC transgenic mice were immunostained for HA (green) and TH (red). Scatterplots show the mean fluorescence per μm process length from individual fields. ****, p<0.0001; ns, not significant by Mann-Whitney test (n=21 fields from three mice for each condition). Scale bar, 5 μm. Error bars indicate SEM.

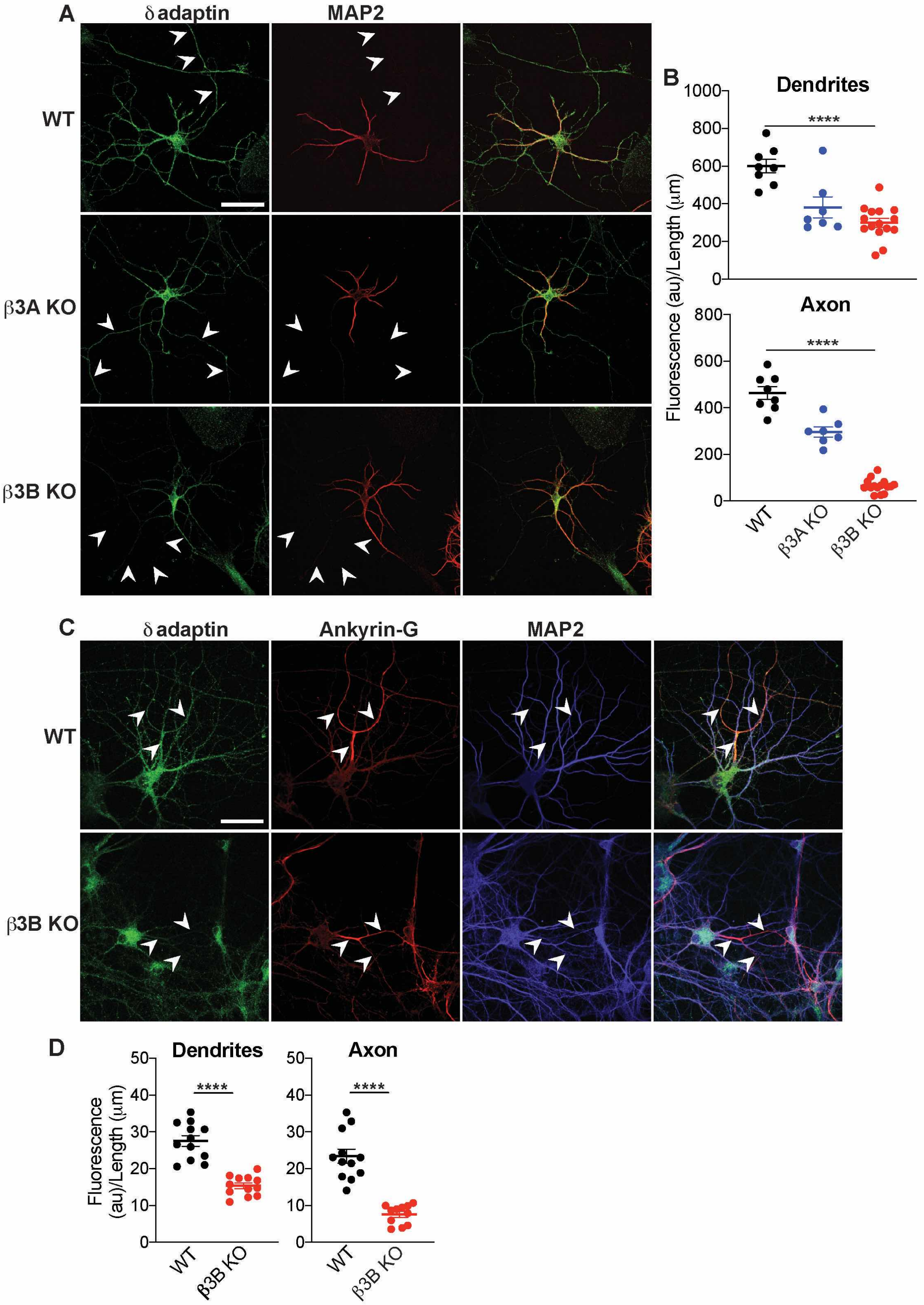

A
δ adaptin
MAP2
WT
β3A KO
β3B KO
B
Dendrites
1000
800
600
400
200
0
****
Axon
800
600
400
200
0
****
Fluorescence (au)/Length (μm)
WT
β3A KO
β3B KO
C
δ adaptin
Ankyrin-G
MAP2
WT
β3B KO
D
Dendrites
Axon
50
40
30
20
10
0
****
Fluorescence (au)/Length (μm)
WT
β3B KO

**Fig. S2. The ubiquitous and neural isoforms of AP-3 differ in localization to the axon.**

(A) Hippocampal neurons from WT, β3A and β3B KO mice were immunostained for δ adaptin (green) and MAP2 (red). Arrowheads indicate the $MAP2^{-}$ axon. Scale bar, 50 µm.

(B) Quantitation of δ adaptin fluorescence per µm process length shows that whereas loss of both isoforms impairs dendritic localization of AP-3, loss of β3B has a larger effect than loss of β3A on axonal localization. **, p<0.01; ****, p<0.0001 by one-way ANOVA with post hoc Bonferroni's test (n=8 WT, 7 β3A KO, 15 for β3B KO). (C) WT and β3B KO hippocampal neurons were stained for δ adaptin (green), ankyrin-G (red) and MAP2 (blue). Arrowheads indicate the ankyrin-G+/$MAP2^{-}$ axons. Scale bar, 50 µm. (D) Quantitation of δ adaptin fluorescence per µm process length shows that the β3B KO preferentially reduces the AP-3 in axons. ****, p<0.0001; by Mann-Whitney test. n=12 coverslips from 3 different cultures for both WT and β3B KO.

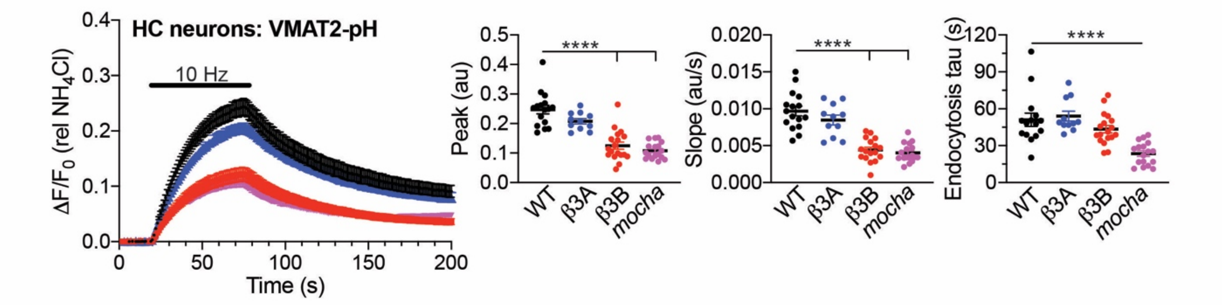


**Fig. S3. Loss of β3B impairs regulated exocytosis and endocytosis of VMAT2 in hippocampal neurons.**

Response of VMAT2-pH to 10 Hz stimulation for 60 s stimulation in hippocampal neurons cultured from WT, β3A, β3B KO and *mocha* mice. Scatterplots show mean peak, mean initial rate of exocytosis and mean endocytic time constant for each coverslip. ****, $p<0.0001$ by one-way ANOVA with *post hoc* Bonferroni's test (n=16 coverslips for WT, 11 for β3A KO, 18 for β3B KO and 17 for *mocha*, each from 3-4 different cultures).

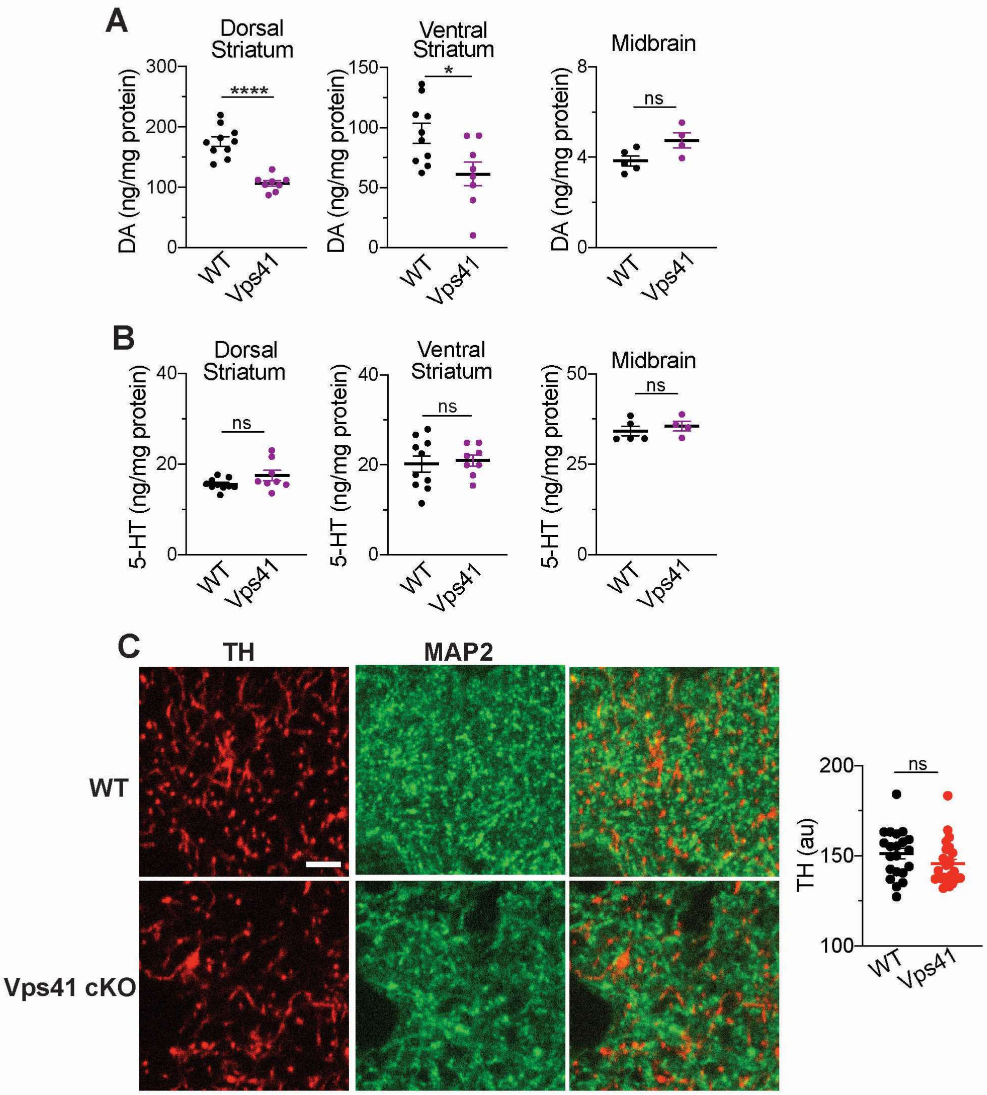


**Fig. S4. Conditional loss of Vps41 reduces axonal dopamine with no change in serotonin (5-HT).**

(A) Tissue content of DA in the dorsal striatum, ventral striatum and midbrain of WT (DATicre+/-) and Vps41cKO mice, presented as scatterplots. ****, p<0.0001; ns non-significant by Mann-Whitney test (n=10 for WT, 8 for Vps41 cKO striatum; n=5 for WT and 4 for Vps41 cKO midbrain).

(B) Tissue content of 5-HT in the dorsal striatum, ventral striatum and midbrain of WT and Vps41 cKO mice, presented as scatterplot. ns, not significant by Mann-Whitney test. n=10 for WT and 8 for Vps41 cKO striatum (n=5 for WT and 4 for Vps41 cKO midbrain).

(C) Striatal sections from WT and Vps41 cKO mice were immunostained for TH (green) and MAP2 (red) and the mean fluorescence per µm process length of individual fields is shown in the scatterplot (right). Scale bar, 5 µm.  ns, not significant by Mann-Whitney (n=? for WT, ? for Vps41 cKO).

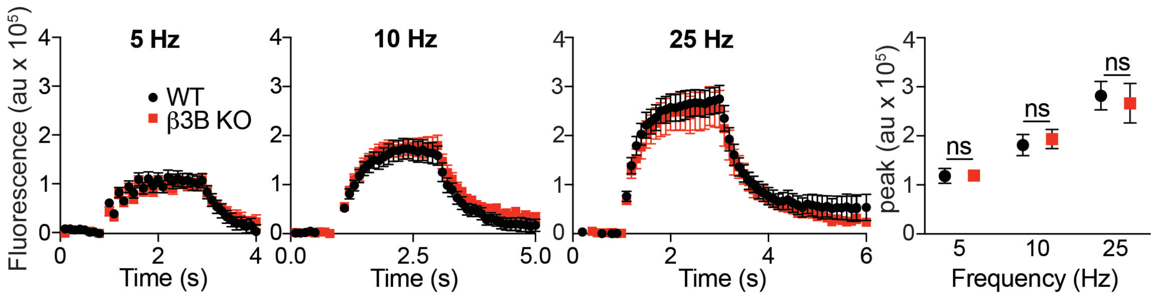


**Fig. S5. Loss of β3B does not affect $Ca^{++}$ entry.**

Midbrain dopamine neurons were loaded with Fluo 5F-AM (2.5 µM) for 15 min, stimulated at different frequencies for 2 s and the peak fluorescence plotted to the right. n=8 coverslips from two independent experiments.   ns, not significant by two-way ANOVA with Sidak's multiple comparison test

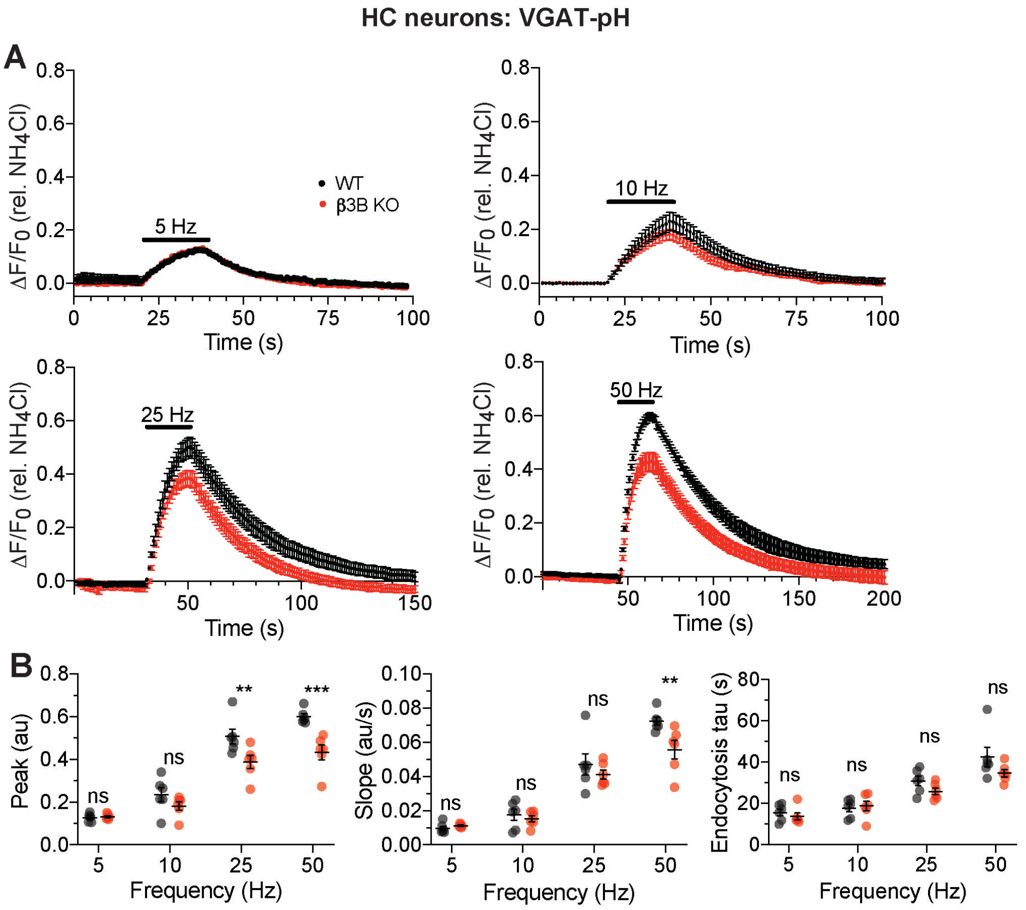


**Fig. S6. Loss of neural AP-3 affects VGAT exocytosis at higher frequency of stimulation**.

(A) Response of VGAT-pH expressed in WT (black) and β3B KO (red) hippocampal neurons to stimulation for 20 s at 5, 10, 25 or 50 Hz. (B) Scatterplots of the mean peak, initial exocytosis slope and endocytosis time constant for WT and β3B KO hippocampal neurons stimulated at different frequencies. ***, $p<0.001$, **, $p<0.01$ by two-way ANOVA With Sidak's *post hoc* test. n=6 coverslips for WT/β3B KO from 2 different hippocampal cultures.

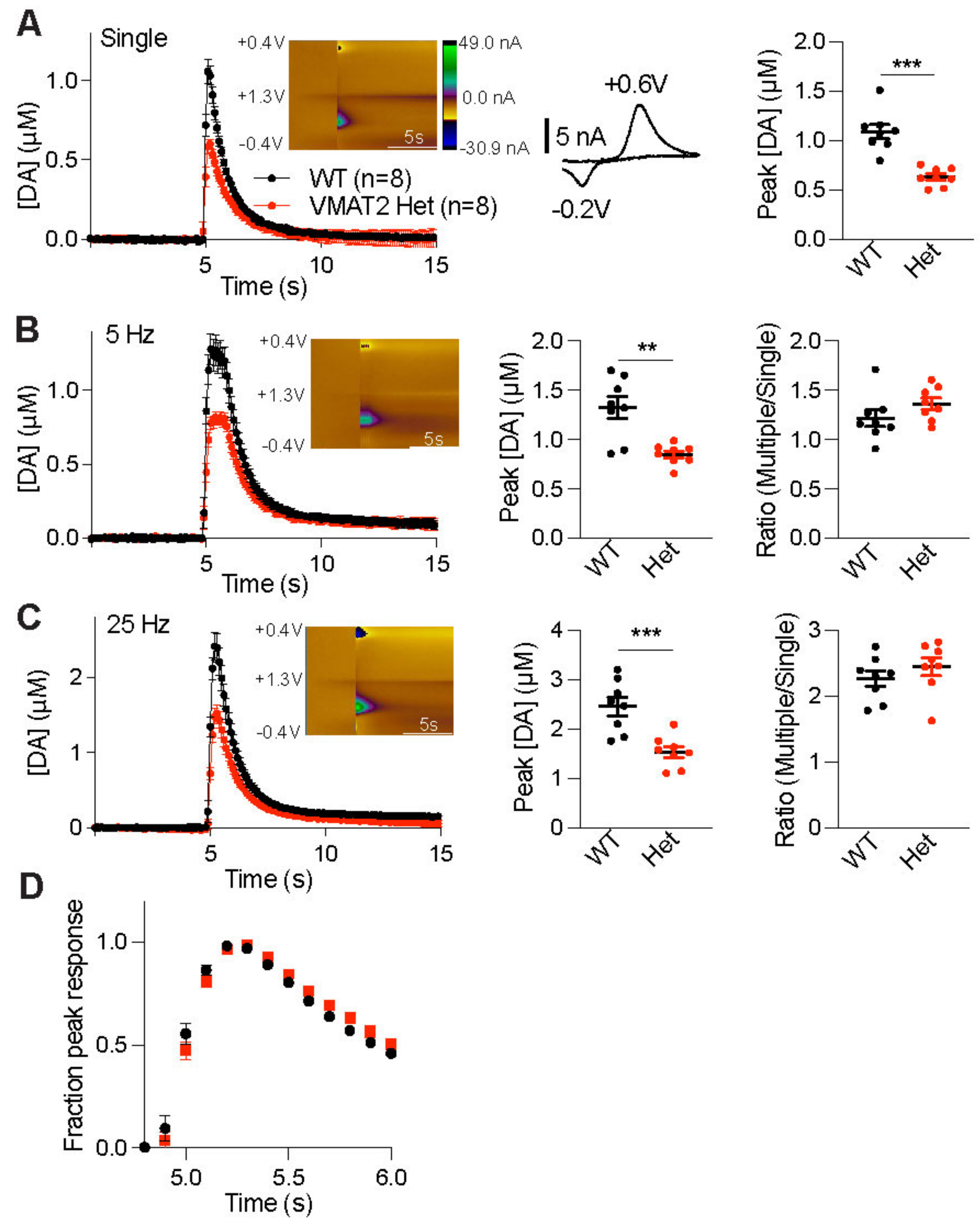


**Fig. S7. Reduction of DA levels does not affect frequency dependence of dopamine release.**

(A) Time course of striatal dopamine release evoked by single stimuli and measured with FSCV in the VMAT2 heterozygote and controls, as described in Fig. 4. Inset: color plot (left). The characteristic dopamine voltammogram (right) shows oxidation and reduction peaks at +0.6 V and -0.2 V, respectively. Scatterplot (right) shows peak dopamine concentrations in VMAT2 Het (n=8 from 4 animals) and WT littermates (n=8 from 4 animals). ***, p<0.001 by Mann-Whitney test (B) Time course of dopamine release evoked by 5 stimuli delivered at 5 Hz. Scatterplots (right) show peak dopamine concentrations (left) and ratio of 5 Hz/single stimulus (right). **, p<0.01. by Mann-Whitney test (C,D) Time course of dopamine release evoked by 5 stimuli delivered at 25 Hz. Scatterplots (right) indicate peak dopamine concentrations (left) and ratio of 25 Hz/single (right). Normalized to the peak response, residual release shows no difference in kinetics between VMAT2 heterozygotes and WT (D). *** p<0.001 by Mann-Whitney test. Error bars indicate SEM.

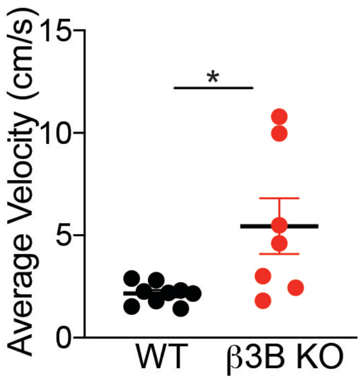


**Fig. S8. Loss of β3B increases movement velocity.**

The spontaneous movement of adult β3B KO (n=7) and WT (n=9) mice was assessed in the open field for 30 minutes. Each dot represents one mouse. Data displayed as mean ± SEM. *, p<0.05 by Mann-Whitney test
